# Preventing escape and malfunction of recoded cells due to tRNA base changes

**DOI:** 10.1101/2024.07.18.604179

**Authors:** Anush Chiappino-Pepe, Felix Radford, Bogdan Budnik, Hüseyin Taş, Teresa L Augustin, Hana M Burgess, Michaël Moret, Azim M Dharani, Qinmei Zheng, Weicheng Fan, Maksud M Africawala, Shova Thapa, Erkin Kuru, Kamesh Narasimhan, Jorge A Marchand, Ramiro M Perrotta, Jonathan M Stokes, Jeantine E Lunshof, John D Aach, Jenny M Tam, George M Church

**Author notes:** Corresponding authors’. ^#^These authors contributed equally to this work (Felix Radford, Bogdan Budnik).

## Abstract

Engineering the genetic code restricts DNA transfer (cellular bioisolation) and enables new chemistries via non-standard amino acid incorporation. These distinct properties make recoded cells state-of-the-art safe technologies. However, evolutionary pressures may endanger the longevity of the recoding. Here, we reveal that recoded *Escherichia coli* lacking 18,214 serine codons and two tRNA^Ser^ can express wild-type antibiotic resistance genes and escape up to seven orders of magnitude faster than expected. We show a two-step escape process whereby recoded cells mistranslate antibiotic resistance genes to survive until modified or mutated tRNAs reintroduce serine into unassigned codons. We developed genetic-code-sensitive kill switches that sense serine incorporation and prevent cellular escape while preserving encoding of three distinct non-standard amino acids. This work lays the foundation for the long-term controlled function of cells that incorporate new chemistries, with implications for the design, use, and biosafety of synthetic genomes in clinical and environmental applications where physical containment is insufficient.

## Introduction

The genetic code defines the rules that link codons, tRNAs, aminoacyl-tRNA synthetases, and amino acids to transfer information from the DNA into proteins. When these rules are known, as they are for *Escherichia coli* (*E. coli*) (Supplementary Fig. 1), organisms can be designed with synthetic, recoded genomes containing a reduced number of codons^1–3^ (Supplementary table 1), and the corresponding release factor^1,4^ and tRNAs^4^ may be deleted. An example is the recoded *E. coli* genome of Ec_Syn61Δ3 that lacks all 18,214 annotated instances of two of the six naturally occurring serine codons (TCA and TCG^2^; or together TCR) and the corresponding tRNA^Ser^ genes (*serT* and *serU*)^4^. Ec_Syn61Δ3 also lacks the amber stop codon^2^ and release factor 1^4^ from its genome. Such free codons have been extensively used for genetically encoding non-standard amino acids^5–7^ with broad, relevant applications, like the discovery of new-to-nature enzyme biochemistries^8^ and the production of peptide drugs^9,10^.

Controlled usage of living technologies for clinical^11–14^ and environmental^15–17^ applications requires measures that prevent cells from escaping designed programs and any engineered functionality from being released into nature. Likewise, such measures may also prevent cells from acquiring mutations and undesired functionalities from surrounding natural environments that promote cellular escape under evolutionary pressures. To prevent cellular escape, several biocontainment strategies have been established over the last 15 years^18^, including engineered nutrient dependencies^19^, altered genetic codes^1,4,20,21^, and kill switches^22–24^. The most stringent biocontainment strategy to date involves the engineering of three fitness relevant enzymes in a recoded cell to depend on a synthetic non-standard amino acid called L4’4’-biphenylanaline (bipA)^19,25^. Unfortunately, such an engineered nutrient dependency is insufficient to prevent escape if recoded cells can receive exogenous DNA through horizontal gene transfer. The altered genetic code and absence of some tRNAs in recoded cells serve as inherent barrier against incoming DNA from retaining functionality, thereby defining a *genetic-code-based cellular bioisolation*. Indeed, it was recently shown that Ec_Syn61Δ3 can be infected with viruses that encode their own tRNA^Ser^ ^20,21^. Repurposing of serine TCR codons to incorporate leucine or other standard amino acids in Ec_Syn61Δ3 prevented virus infection^20,21^. But, by allocating free codons to other natural amino acids, we miss the opportunity to use them for non-standard amino acids and thus miss one of the key benefits of genomic recoding.

Here, we discovered another vulnerability of a genetic-code-based cellular bioisolation that does not require the transferred DNA to code for tRNAs or be optimally expressed for immediate functionality, such as the case with viruses that carry their own translation machinery^26,27^. We show that Ec_Syn61Δ3 can express horizontally transferred, TCR-containing antibiotic resistance genes that cause eventual escape from the genetic-code-based cellular bioisolation. We identified a two-step mechanism by which recoded cells acquire antibiotic resistance through gene transfer. First, recoded cells mistranslate antibiotic resistance genes, and then available tRNA^Ser^ undergoes modifications or mutations to re-incorporate serine into TCR codons. To prevent the expression of antibiotic resistance genes with TCR codons, we successfully engineered genetic-code-sensitive kill switches that reduced the escape rate up to four orders of magnitude to 10^-8^ cfu/mL upon antibiotic selection. Our genetic-code-sensitive kill switches allow full retention of the ability to incorporate one to three distinct non-standard amino acids in Ec_Syn61Δ3.

This work showcases how evolutionary pressures can shape an engineered genetic code^28^ and lays the foundation for safely using cells that can genetically encode multiple non-standard amino acids. We discuss the need for adaptive consensus guidelines on biosafety and biosecurity that can account for future discoveries in genome design^29^. This study has broad implications for the design, application, and biosafety of synthetic genomes for clinical and environmental applications in which physical containment is not sufficient.

## Results

### Antibiotic resistance genes promote escape from a genetic-code-based cellular bioisolation

The recoded genome of Ec_Syn61Δ3 lacks all annotated instances of TCR serine codons and the cognate tRNA^Ser^ genes *serT* and *serU.* This fact led us to expect that Ec_Syn61Δ3 supplied with a plasmid encoding a wild-type TCR-codon-containing antibiotic resistance gene would be susceptible to the antibiotic. However, experiments comparing a high-copy plasmid pUC57 expressing a beta-lactamase gene with three TCR codons (two TCA and one TCG, Fig. 1a, Supplementary table 2) with a recoded TCR-less version of the same plasmid (pUC57-dTCR) showed the opposite. We found that transformation of the wild-type plasmid into Ec_Syn61Δ3 followed by selection on carbenicillin-containing agar plates at standard concentration for 2-3 days yielded hundreds to thousands of colony forming units per mL (cfu/mL). We measured a frequency of ∼1.3x10^-4^ cfu/mL (*n = 4*) Ec_Syn61Δ3 escapees with wild-type pUC57 not susceptible to antibiotics (Fig. 1b, Methods). An expected antibiotic resistance acquisition rate through spontaneous genomic nucleotide mutations in *E. coli* is in the order of 10^-9^ or 10^-10^ cfu/mL^30^. Thus, the observed escape rate of Ec_Syn61Δ3 with wild-type pUC57 is around five orders of magnitude higher than expected. We verified that the plasmid-less strain Ec_Syn61Δ3 does not acquire carbenicillin resistance at a rate higher than 10^-9^ cfu/mL (Fig. 1b, Supplementary Fig. 2), which suggested that the plasmid pUC57 with a TCR-containing beta-lactamase gene conferred a fitness advantage to Ec_Syn61Δ3 under carbenicillin selection.

**Fig. 1.**
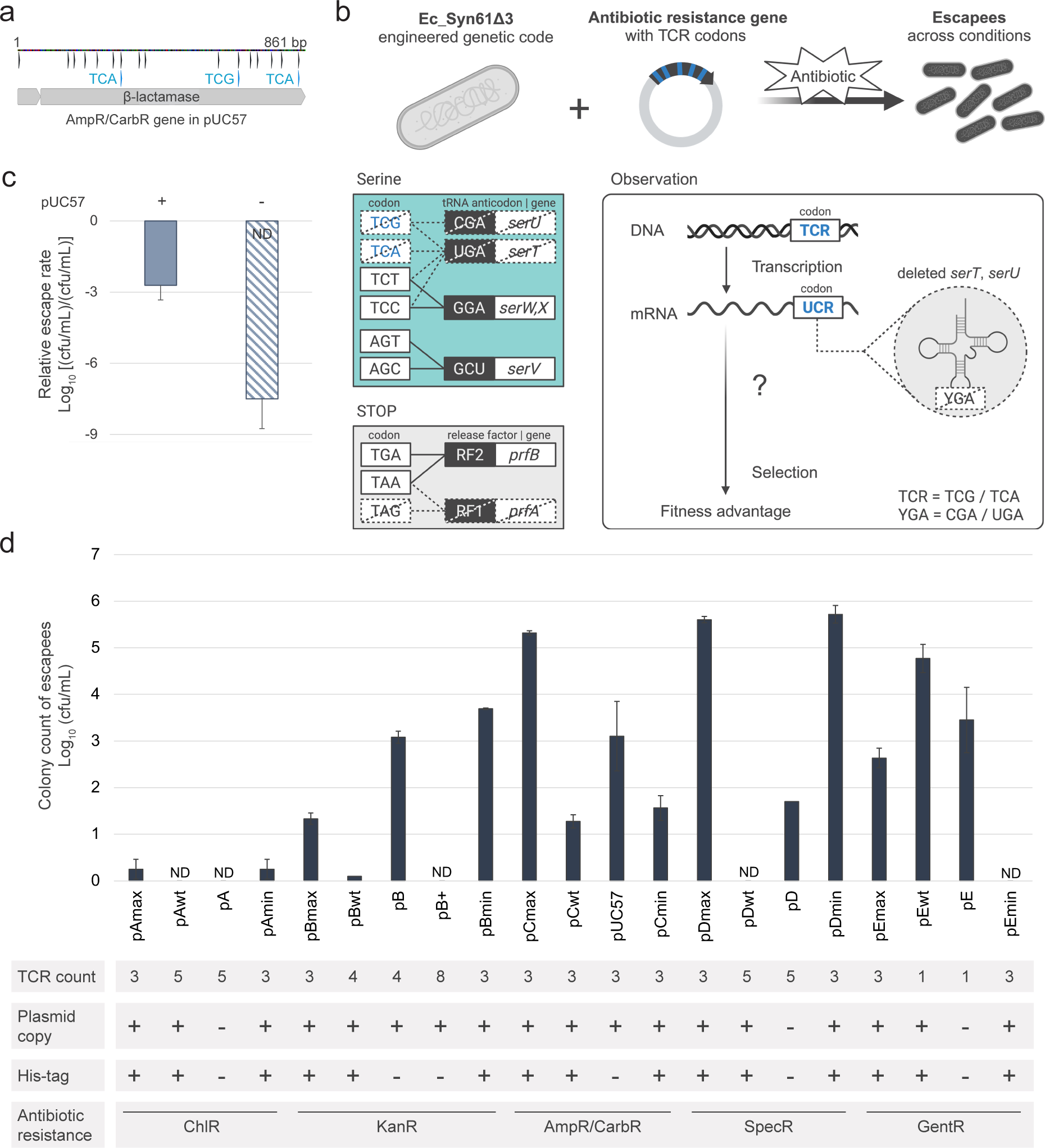
Escape rate of Ec_Syn61Δ3 with TCR-containing antibiotic resistance genes. **a**, Wild-type beta-lactamase gene in plasmid pUC57, conferring ampicillin/carbenicillin resistance. Black arrows locate all wild-type serine codons, and blue arrows highlight TCA/TCG (TCR) codons. **b**, Top: Schematic representation of the transformation and selection process performed with all plasmids to study the escape rate of Ec_Syn61Δ3. Bottom left: Engineered genetic code of Ec_Syn61Δ3. Missing codons, anticodons, and genes are marked with dashed lines. Remaining codons, anticodons, and genes are marked with bold lines. Bottom right: Schema of central dogma of biology including our observation. **c**, Relative escape rate [(cfu/mL)/(cfu/mL)] defined as the number of colonies formed by Ec_Syn61Δ3 with (+) and without (-) the wild-type plasmid pUC57 relative to those of parallel experiments with pUC57-dTCR upon selection onto carbenicillin-containing agar plates (*n = 4*). Escapees of Ec_Syn61Δ3 without the pUC57 plasmid were not detected (ND) onto carbenicillin-containing agar plates with up to 1 mL of OD_600_ plated (Methods). The dashed bar length represents the ratio of 10^-9^ cfu/mL and measured number of cells of Ec_Syn61Δ3 with pUC57-dTCR (*n = 4*). **d**, Number of colonies formed by Ec_Syn61Δ3 escapees with selected plasmids including TCR-containing antibiotic resistance genes upon selection onto antibiotic-containing agar plates (mean ± standard deviation (SD), *n = 2-4*). The table below describes the number of TCR codons, plasmid copy number, presence of a his-tag on the antibiotic resistance protein, and antibiotic resistance for each plasmid. High or low plasmid copy number is denoted by + or -. Presence or absence of a His-tag is denoted by + or -. Antibiotic resistance acronyms: ChlR, chloramphenicol; KanR, kanamycin; AmpR/CarbR, ampicillin/carbenicillin; SpecR, spectinomycin; and GentR, gentamicin. Designs tagged as “max” or “min” contain TCR codons in serine positions with the highest or lowest probability for serine, respectively, based on the protein language model ESM-1b (Supplementary Fig. 3). Antibiotic acronyms: A, chloramphenicol; B, kanamycin; C, carbenicillin; D, spectinomycin; and E, gentamicin.

To evaluate the breadth of antibiotic selection scenarios in which Ec_Syn61Δ3 could escape and the influence of TCR-containing gene designs on the escape rate, we expanded the set of plasmids to include six readily available and fifteen new designs (Supplementary table 2). We hypothesized that the escape rate would be affected by the positions of TCR codons and the probability that serine is important for protein fitness (essential) in those TCR positions. Machine learning models trained with millions of protein sequences (protein language models) learn the contextual importance of an amino acid in a sequence^31^. Hence, we used a protein language model (Evolutionary Scale Model, ESM-1b^32^) to design antibiotic resistance genes with TCR codons in wild-type serine positions in which serine would likely be the most or least essential for protein function (Methods, Supplementary Fig. 3). Our plasmid set included 21 genes in low and high copy number plasmids, tagged or not with a poly-histidine tail, and genes conferring antibiotic resistance to chloramphenicol, kanamycin, carbenicillin, spectinomycin, and gentamicin with one to eight TCR codons (Supplementary Dataset 1).

Astonishingly, we observed escapees for all antibiotic selections at standard concentrations when the antibiotic resistance genes contained up to five TCR codons (Fig. 1d). The frequency of escapees, measured as the ratio of cfu/mL formed in antibiotic selection over the cfu/mL formed without antibiotic selection after transformation, was as high as 10^-3^ cfu/mL (Fig. 1d). We verified that our designed plasmids rendered comparable numbers of colonies in an *E. coli* strain with tRNA^Ser^ genes *serT* and *serU*, which confirmed that all antibiotic resistance gene designs were functional and expressed similarly in the presence of cognate tRNA^Ser^ serT and serU (Supplementary Fig. 4). We also confirmed that the same cell concentration of the plasmid-less Ec_Syn61Δ3 strain did not lead to escape in all our selection conditions (Supplementary Fig. 2).

Our results confirmed that a broad set of TCR-containing antibiotic resistance genes conferred a fitness advantage to Ec_Syn61Δ3 in the presence of antibiotics, highlighting a generalizable, previously unknown weakness in a genetic-code-based cellular bioisolation. Our gene designs from the protein language model ESM-1b (Supplementary Fig. 3) with the same number of TCR codons in distinct positions would result in up to four orders of magnitude difference in escape rate. Both the number and positions of TCR codons, derived from ESM-1b, influenced Ec_Syn61Δ3’s escape, supporting our hypothesis that the escape rate is affected by the probability that serine is essential in specific (TCR) positions for protein fitness. To provide quantitative guidance in the selection of TCR-containing antibiotic resistance genes that minimize Ec_Syn61Δ3 escape, we derived an equation that uses amino acid probabilities from protein language models (Methods). These results motivated us to further explore the mechanism of Ec_Syn61Δ3 escape with TCR-containing antibiotic resistance genes.

### Translation error and tRNA^Ser^ base changes enable recoded cells to escape

We next investigated the mechanism of escape of Ec_Syn61Δ3 upon antibiotic selection in the presence of TCR-containing antibiotic resistance genes. For this purpose, we collected genome and plasmid sequences for over a hundred escapees. We also collected proteomics, tRNA-seq, and dynamic fluorescence data based on a newly designed reporter to analyze the escape process (Fig. 2a, Supplementary table 3).

**Fig. 2.**
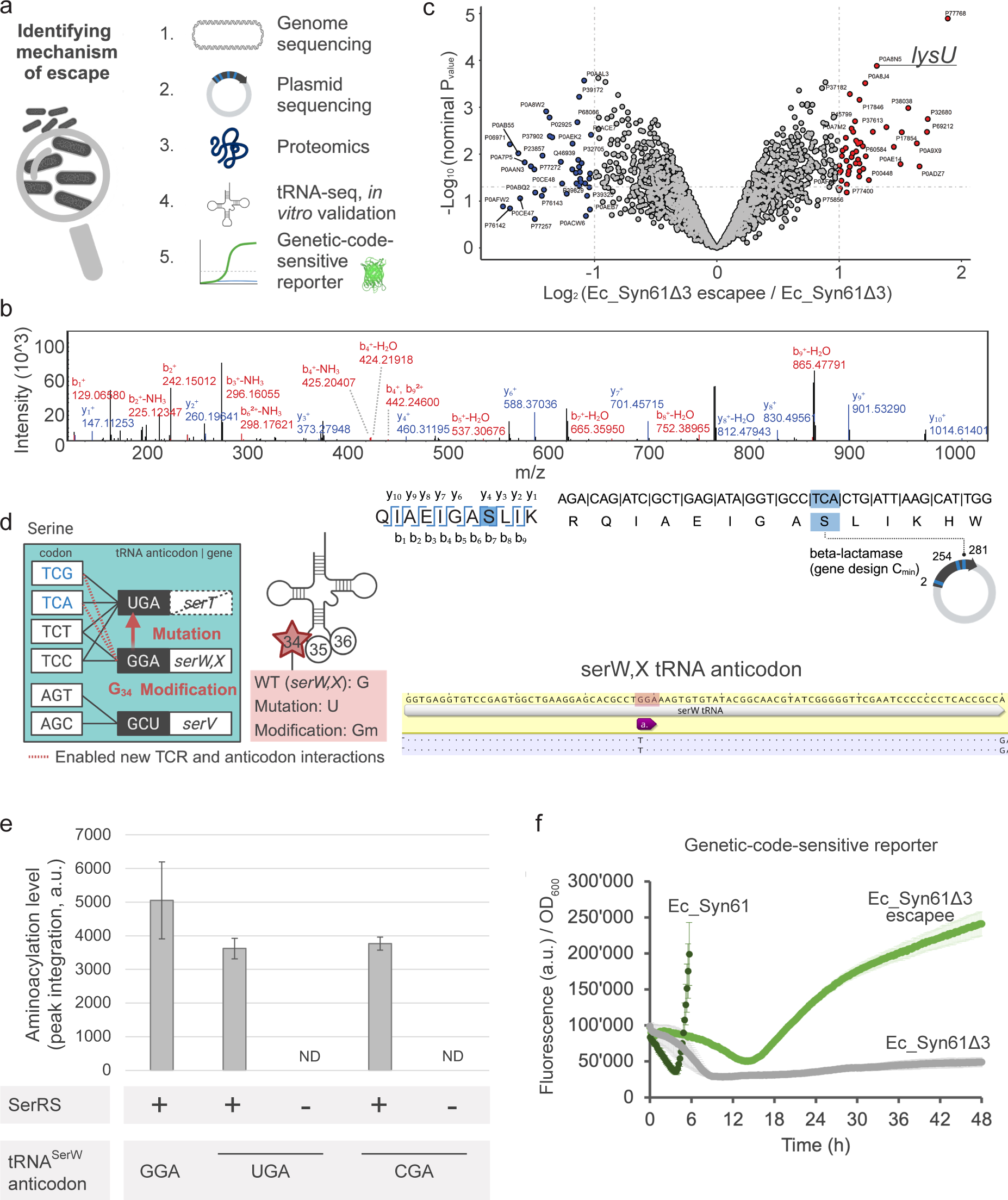
Quantification of mistranslation and tRNA^Ser^ base changes in escapees of Ec_Syn61Δ3. **a**, Schematic representation of data collected to identify the mechanism of escape in Ec_Syn61Δ3. **b**, Mass spectrum of the beta-lactamase enzyme (encoded in a gene containing 3 TCR codons, pCmin design) as detected in escapees of Ec_Syn61Δ3. We detected serine incorporated into the third TCR codon with the peptide QIAEIGASLIK (see Supplementary Figures 7-11 for complementary data). **c**, Differential, untargeted proteomics analysis of Ec_Syn61Δ3 escapees with the pUC57 plasmid relative to Ec_Syn61Δ3 (*n = 3*). We highlight the overexpressed *lysU*. **d**, Detection of mutation in the anticodon of the tRNA^Ser^ serW/X G34U. **e**, *In vitro* aminoacylation of the wild-type tRNA^Ser^ serW/X and the mutants of serW/X with anticodons UGA and CGA show similar aminoacylation levels. **f**, Fluorescence per OD_600_ over time in Ec_Syn61Δ3 carbenicillin escapees with our genetic-code-sensitive reporter (light green), compared to fluorescence of this reporter in Ec_Syn61 with tRNA^Ser^ genes *serT* and *serU* (dark green) and in Ec_Syn61Δ3 when it is part of a plasmid with a recoded carbenicillin resistance gene (gray) (mean ± SD; *n = 5*).

Through whole-genome sequencing and amplification of genomic regions, we confirmed that genes *serU* and *serT* of wild-type tRNA^Ser^ cognate to TCR codons were not present in the genome of Ec_Syn61Δ3 escapees (Supplementary Fig. 5). We did not observe known antibiotic resistance mutations in Ec_Syn61Δ3 escapees other than single nucleotide mutations in ribosomal genes, which favor mistranslation^33^ (Supplementary table 4). Separately, plasmid sequence data from Ec_Syn61Δ3 escapees confirmed that all genes kept multiple TCR codons (Supplementary table 5). We also observed an increase in the copy number of antibiotic resistance genes within plasmids (Supplementary Fig. 6).

To evaluate whether TCR-containing, antibiotic resistance genes are expressed in Ec_Syn61Δ3 and to identify amino acids incorporated on TCR codon positions, we purified antibiotic resistance enzymes from Ec_Syn61Δ3 escapees from four gene designs and analyzed them by phase liquid chromatography and tandem mass spectrometry (LC–MS/MS). We detected peptides, indicating protein expression, across the sequences of beta-lactamase (carbenicillin resistance) and aminoglycoside phosphotransferase (kanamycin resistance) for all analyzed samples of Ec_Syn61Δ3 escapees. This result shows that Ec_Syn61Δ3, despite its lack of cognate tRNA^Ser^ genes *serT* and *serU*, can translate genes with three instances of theoretically unreadable TCR codons.

The amino acids being incorporated in TCR positions included serine, proline, arginine, and threonine (Supplementary Fig. 7-10, Supplementary table 5). Interestingly, the lowest incorporation rate of serine occurred in the TCR positions of aminoglycoside phosphotransferase with the lowest serine probability based on a protein language model (Supplementary Fig. 3 and 9). These results suggest that mistranslation initially occurs in Ec_Syn61Δ3 to enable expression of antibiotic resistance enzymes and survival upon antibiotic selection. We could not detect antibiotic resistance enzymes from TCR-containing genes in Ec_Syn61Δ3 without antibiotic selection, which confirms that selection pressure plays a role in the expression rate of TCR-containing genes and escape rate of Ec_Syn61Δ3.

At least 93% of all detected beta-lactamase and aminoglycoside phosphotransferase from four gene designs (*n = 9*) contained serine in TCR positions (Fig. 2b, Supplementary Fig. 7-10), indicating that the incorporation of serine in TCR codons is favored compared to other amino acids. To evaluate whether Ec_Syn61Δ3 escapees recovered their ability to incorporate serine in TCR codons through a new mechanism of escape that affected also non-essential genes, we designed a novel genetic-code-sensitive reporter, consisting of a GFP coding gene with TCA codons in five serine positions with the highest probability of being essential for function based on ESM-1b^32^. LC–MS/MS analysis of the genetic-code-sensitive GFP reporter verified that serine was being incorporated in instances of TCA codons of GFP with traces (<8%) of proline (Supplementary Fig. 11, *n = 9*). The high (>92%) incorporation rate of serine in our genetic-code-sensitive reporter, which does not confer any fitness advantage to escapees, strongly suggests that Ec_Syn61Δ3 escapees recovered the ability to incorporate serine in TCR codons.

We next sought to understand the mechanism by which Ec_Syn61Δ3 escapees can incorporate back serine into TCR codons. Untargeted deep proteome analysis showed an altered proteome in escapees with the pUC57 plasmid upon carbenicillin selection relative to the controls Ec_Syn61Δ3 and two parental *E. coli* strains with *serT* and *serU*, *i.e.*, Ec_Syn61 and MDS42 (Fig. 2c). We observed overexpression of a promiscuous lysine aminoacyl tRNA synthetase (lysU|P0A8N5, fold change (FC) = 2.37, *n = 3,* Fig. 2b). Interestingly, lysU has been shown to catalyze misacylation of tRNA^Lys^ with serine, arginine, and threonine^34,35^, which we detected in antibiotic resistance enzymes and GFP of Ec_Syn61Δ3 escapees (Supplementary Fig. 7-11). Four enzymes that modify tRNAs were also slightly overexpressed (1.3<FC<2). For example, tRNA m^5^U54 methyltransferase (trmA|P23003, FC = 1.42, *n = 3*) and an uncharacterized tRNA methyltransferase (lasT|P37005, FC = 1.39, *n = 3*). tRNA modifications are known to expand the repertoire of codon-anticodon wobble interactions^36,37^ and alleviate ribosome stalling^38^. In alternative genetic codes, methylation at N-7 of wobbling guanosine endows the tRNA with the capability of forming base pairs with all four nucleotides, A, U, G, and C^39,40^. Available tRNA^Ser^ in Ec_Syn61Δ3, like serW/X with GGA anticodon might recognize TCR codons should they undergo methylation at the wobbling guanosine by the overexpressed tRNA modifying enzymes (trmA|P23003 or lasT|P37005). Based on these observations, we hypothesize that mistranslation and modification of tRNAs are involved in the escape of Ec_Syn61Δ3.

Through tRNA-seq, we next looked for tRNAs able to incorporate serine in TCR codons of Ec_Syn61Δ3 escapees. We first investigated tRNA^Ser^ from genes present in the Ec_Syn61Δ3 genome like *serW/X*, which should not recognize TCR codons either through Watson-Crick or wobble interactions in its wild-type form (Supplementary Fig. 1). Surprisingly, tRNA-seq data showed that a fraction (1-5%) of available tRNA^Ser^ serW/X from our population of escapees presented a single point mutation at position 34. The mutation G34U lies in the anticodon GGA of serW/X and reintroduced the anticodon UGA and, with it, the ability to decode TCR codons (Fig. 2d). Through an *in vitro* aminoacylation reaction, we confirmed the ability of serW/X mutants (mutated to include a UGA or CGA anticodon) to be aminoacylated with serine by purified *E. coli* seryl-tRNA synthetase to an average 73±1% level of the wild-type serW/X (Fig. 2e), consistent with the understanding that seryl-tRNA synthetase does not recognize the tRNA^Ser^ anticodon for aminoacylation^41^. These results suggest that after an initial mistranslation stage, Ec_Syn61Δ3 can eventually escape antibiotic selection by mutating or modifying available tRNAs to render functional mutant tRNA^Ser^. Such mutants tRNA^Ser^ allow Ec_Syn61Δ3 to recognize TCR codons for serine incorporation and thereby express genes with a standard genetic code (Supplementary Fig. 7-11).

To track the modification and mutation rate of tRNA^Ser^ *in vivo* for incorporation of serine, we used our genetic-code-sensitive GFP reporter (Supplementary Fig. 11), which allowed us to dynamically measure fluorescence as a function of growth and time at high throughput, thereby suggesting a time point at which Ec_Syn61Δ3 escapees can incorporate serine back into TCR codons. Starting at the mid-exponential phase, we measured a ∼5-fold increase in fluorescence per cell density (OD_600_) over time in escapees of Ec_Syn61Δ3 under carbenicillin selection (Fig. 2f). For reference, we observed an increase rate of fluorescence over OD_600_ at a 14.5-fold faster rate from the beginning of the exponential phase in an *E. coli* strain with tRNA^Ser^ genes *serT* and *serU* (Ec_Syn61) compared to those observed in escapees of Ec_Syn61Δ3 (Supplementary Fig. 11). Our genetic-code-sensitive reporter and other reporter designs with more than 2 TCR codons did not yield fluorescence in Ec_Syn61Δ3 when they were part of a plasmid with a recoded antibiotic resistance gene (Fig. 2f, Supplementary Fig. 12, Dataset 2). Based on these results, we hypothesize that the lack of a TCR-containing gene conferring fitness advantage does not evolve recoded cells to enable expression of TCR-containing genes, as is the case when there exists a TCR-containing antibiotic resistance gene (Fig. 2f).

The observations with our genetic-code-sensitive reporter (Supplementary table 7) are in accordance with our separate sequencing and omics data for antibiotic resistant escapees (Supplementary tables 4-6) and validate our live reporting approach for the escape of Ec_Syn61Δ3 through tRNA^Ser^ modifications or mutations (hereafter referred as tRNA^Ser^ base changes).

### Design and implementation of genetic-code-sensitive kill switches

To prevent the escape of a recoded cell *via* tRNA^Ser^ base changes, we next designed and implemented genetic-code-sensitive kill switches that penalized the incorporation of serine into TCR codons while simultaneously enabling the incorporation of multiple non-standard amino acids in Ec_Syn61Δ3. Our kill switches contain TCR codons at selected essential serine positions of antibacterial proteins (Methods) to satisfy three properties. First, the antibacterial gene is not translated when there is no cognate tRNA^Ser^. Second, the antibacterial gene is nonfunctional when there is a cognate tRNA incorporating a non-standard amino acid. Lastly, the antibacterial gene is expressed and toxic when a tRNA^Ser^ enables the incorporation of serine into TCR codons (Fig. 3a).

**Fig. 3.**
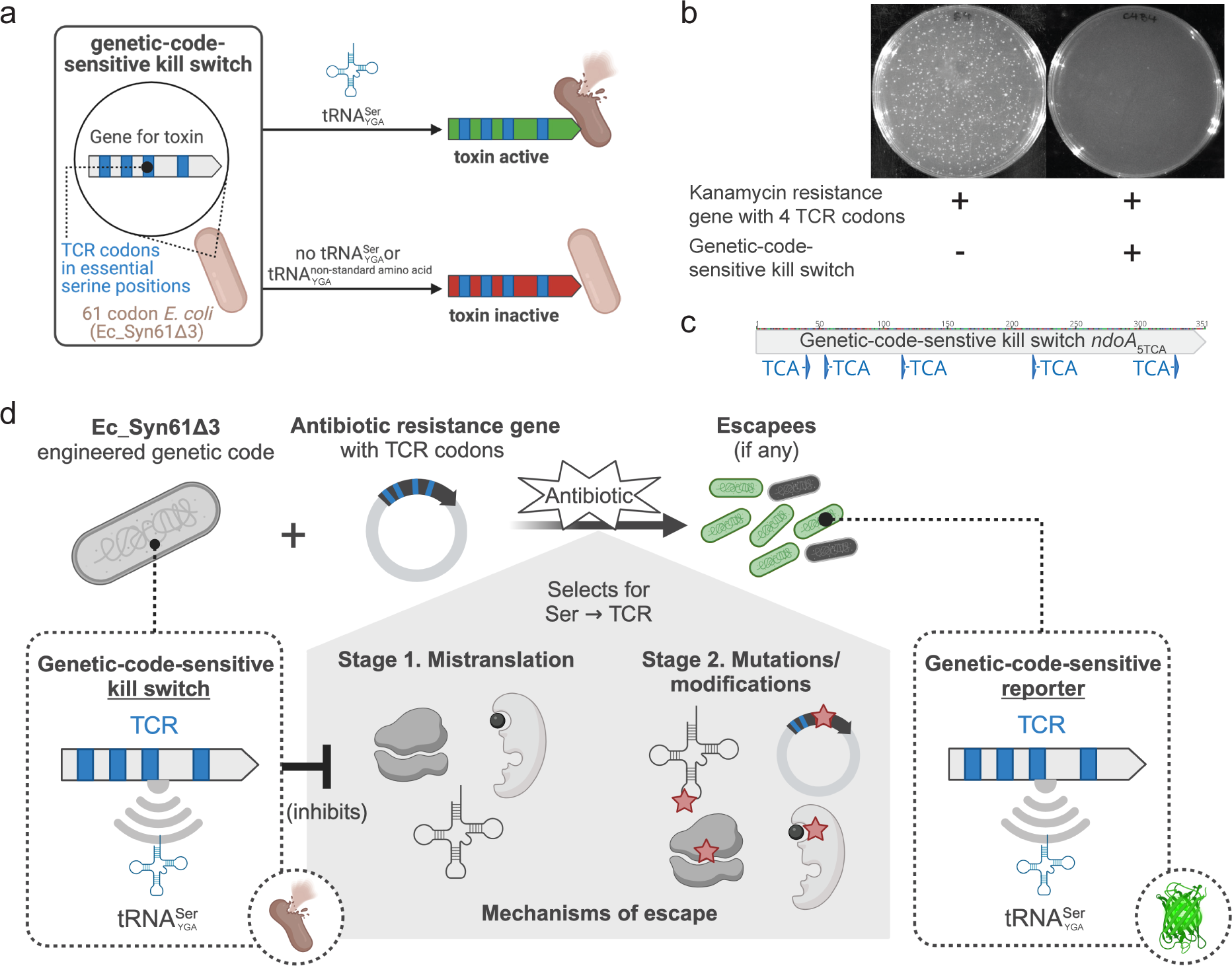
Design and performance of genetic-code-sensitive kill switches in Ec_Syn61Δ3. **a**, Conceptual representation of the on/off status of a genetic-code-sensitive kill switch in Ec_Syn61Δ3. **b**, Escape rate of Ec_Syn61Δ3 without (left) and with (right) a successful genetic-code-sensitive kill switch design in the presence of a kanamycin resistance gene with four TCR codons (*n = 3*). **c**, A successful genetic-code-sensitive kill switch is the antibacterial *ndoA* with five instances of TCA codons in wild-type serine positions. **d**, Our genetic-code-sensitive kill switches stop escape through expression of TCR-containing antibiotic resistance genes for a set of alternative (observed and hypothetical) mechanisms of escape. If there are escapees from antibiotic resistance genes through tRNA^Ser^ modifications and mutations and from our genetic-code-sensitive kill switches, our genetic-code-sensitive reporter will highlight these. In this figure, tRNA^Ser^ refers to serine tRNAs with YGA anticodon that recognize TCR codons. We represent the genetic-code-sensitive gene expression as a radar for tRNA^Ser^ recognizing TCR codons.

We selected a diverse set of antibacterials from bacterial toxin-antitoxin systems^42–45^ (Supplementary Fig. 13) for our genetic-code-sensitive kill switches (Supplementary table 8). For these designs we conducted experiments confirming that our three desired properties were satisfied: (1) the presence of TCR-containing antibacterial genes alone in Ec_Syn61Δ3 did not affect growth; (2) the simultaneous presence of TCR-containing antibacterial genes and an orthogonal translation machinery incorporating the non-standard amino acid bipA also did not affect growth; (3) the expression of a TCR-recognizing tRNA^Ser^ serT or serU yielded severe growth reduction or cell death (Supplementary Fig. 14).

The ultimate evaluation of our genetic-code-sensitive antibacterials required testing them in Ec_Syn61Δ3 and in the presence of TCR-containing antibiotic resistance genes known to yield a high escape rate (Fig. 1c). The Ec_Syn61Δ3 escape rate varied in response to alternative antibacterial proteins, in agreement with previous literature^22,46^, and to the number of TCR codons in our gene designs (Supplementary Fig. 15). Importantly, we found a successful genetic-code-sensitive kill switch with the antibacterial *ndoA* containing five instances of TCA codons (Fig. 3c). This kill switch reduced the escape rate of Ec_Syn61Δ3 four orders of magnitude to 10^-8^ cfu/mL compared to the thousands of colonies formed by the same concentration of Ec_Syn61Δ3 cells without the genetic-code-sensitive kill switch and in the presence of a kanamycin resistance gene containing four TCR codons (Fig. 3b). These results further verified our findings on the mechanism of escape by Ec_Syn61Δ3 due to tRNA^Ser^ base changes and validated our genetic-code-sensitive kill switches (Fig. 3d).

## Discussion

This work showcases how evolutionary pressures reshape an engineered genetic code^28^. It also advances prior methods for bio-containing genomes by identifying new needs and means for preventing escape. We demonstrate that 18,214 recoded codons and two deleted tRNA^Ser^ genes in the Ec_Syn61Δ3 synthetic genome are insufficient to restrict the exchange of genetic information even if incoming DNA does not encode tRNA^Ser^ cognate to TCR. We show that the translation machinery, including ribosomes, aminoacyl-tRNA synthetases, tRNA modifying enzymes, and tRNAs, can enable the expression of genes with codon sets for which no cognate wild-type tRNA is present in the initial genome.

Here, we present our hypothesis for a two-step mechanism of escape involving known and newly discovered molecular strategies linked to the central dogma and antibiotic resistance acquisition. First, Ec_Syn61Δ3 increases copy number and mistranslates TCR-containing genes, which agrees with phenotypes seen in bacteria under antibiotic stress^33,47^. Second, we discovered that Ec_Syn61Δ3 restores its ability to incorporate serine into TCR codons. We detected tRNA^Ser^ mutations, moderate overexpression of tRNA modifying enzymes, and overexpression of a promiscuous lysine-tRNA synthetase, which can misincorporate serine^34^ (Supplementary Fig. 16a). The observed mechanism of escape suggests that the genetic code of Ec_Syn61Δ3 and other genetic code schemes, like a serine-leucine swapping^20^, are at risk of escape through modifications of the natural translation machinery.

To our knowledge, the expression of TCR-containing genes in Ec_Syn61Δ3 without viral tRNA^Ser^ had not been reported, possibly because previous studies had not considered genes that might confer a fitness advantage or used genes with large number of TCRs^4,21^, which are not readily expressed in Ec_Syn61Δ3, as we showed (Supplementary Fig. 12). One prior study observed thousands of colonies from Ec_Syn61Δ3 on kanamycin-containing agar plates when Ec_Syn61Δ3 was transformed with a plasmid including both a kanamycin resistance gene with four TCR codons and tRNA^Ser^ variants recognizing TCR codons^20^. It was suggested that those tRNA^Ser^ were used to express the kanamycin resistance gene in Ec_Syn61Δ3. However, the identified tRNA^Ser^ variants could not rescue viral replication, which was puzzling^20^. We used a plasmid with the same kanamycin resistance gene but without tRNA^Ser^ and it yielded thousands of colonies (Fig. 1d), indicating that previously identified tRNA^Ser^ may not be required to express the kanamycin resistance enzyme.

Through expression of antibiotic resistance genes containing TCR codons, we show that Ec_Syn61Δ3 acquired antibiotic resistance at a rate as high as 10^-2^ cfu/mL. Based on this observation, we suggest that recoded genomes might be evolutionarily driven to regain their native standard amino acid recognition potential and express horizontally transferred genes yielding a fitness advantage. In this sense, a recoded cell resembles the unstable state of a bistable system (Supplementary Fig. 16b), whereby reintroducing the capability to incorporate the naturally cognate amino acid in the unassigned codon might stabilize the recoded cell more than enabling or maintaining the reassigned codon to incorporate a non-standard amino acid. Factors like gene relevance for cell fitness, expression level (given by the promoter, ribosome binding site, and plasmid/gene copy number), number of TCR codons, and positions of TCR codons played a relevant role in the probability of TCR-containing gene expression and escape in Ec_Syn61Δ3 (Supplementary Fig. 16c). Through a newly derived formula (Methods) using a protein language model^32^, we provide quantitative guidance for the design of genes that are less likely to be expressed in recoded cells. In the future, machine learning models with codon resolution^48^ will facilitate the design of recoded genes and genomes.

The observed expression of genes with a set of TCR codons in Ec_Syn61Δ3 puts at risk the use of this bacterium in natural environments for therapeutics or bioremediation in which physical containment is not sufficient. TCR codons are rare, and only a few occur in genes with a wild-type genetic code, such as antibiotic resistance genes in mobile genetic elements or phage plasmids that are common in nature^49–51^. The possibility of antibiotic resistance genes partially breaking the bioisolation of a recoded cell is an important consideration because the modified cell would remain broadly resistant to viruses through the recoded amber stop codon^1^ and newly resistant to antibiotics through expression of horizontally transferred genes. Our data indicate that incorporating genetic-code-sensitive kill switches, such as the antibacterial *ndoA* with five instances of TCA codons, can significantly reduce the likelihood of escape through the observed means. Based on these key findings, we recommend their inclusion, together with our genetic-code-sensitive reporter and other biocontainment mechanisms, in biosafety guidelines^29^ for recoded organisms (Fig. 4). Consensus guidelines on biosafety and biosecurity need to be adaptive to account for the dynamics inherent to discoveries in genome design.

**Fig. 4.**
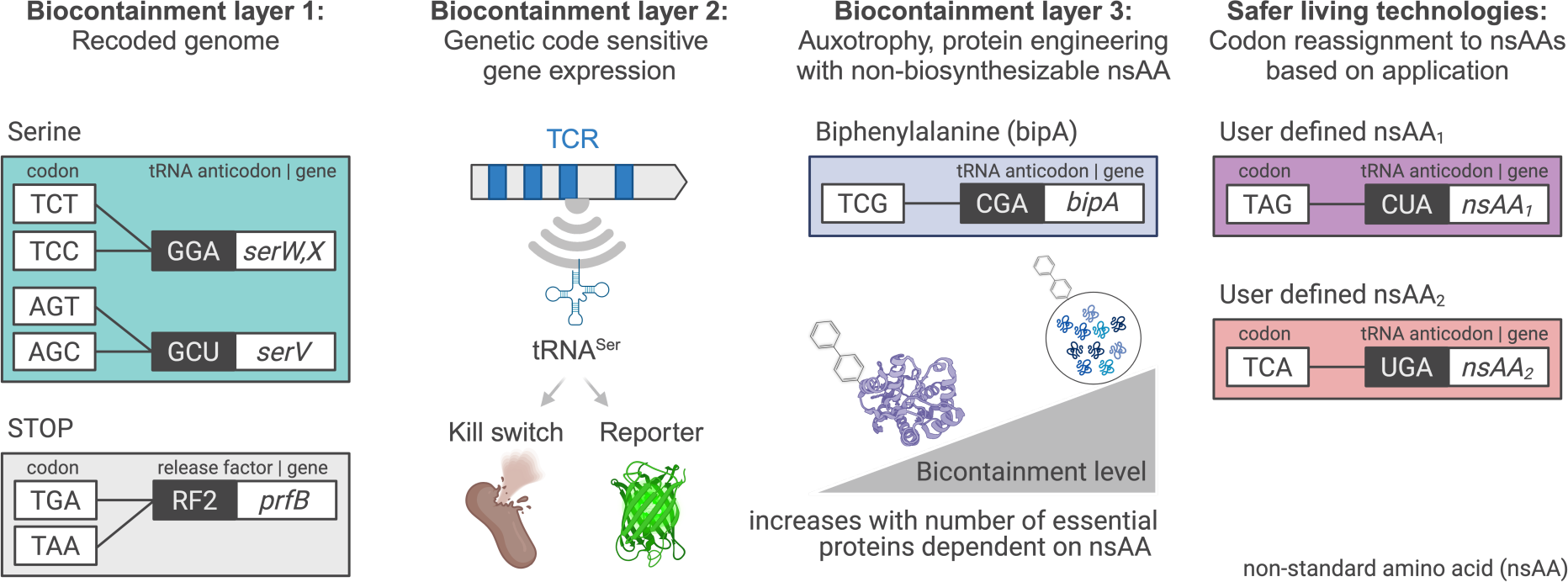
Proposed strategy for the design and application of controlled living technologies. We propose to include at least three layers of biocontainment to develop safer, controlled living technologies. First, genomic recoding will reduce expression of genes, primarily if they are not relevant for fitness and contain more than five TCR/unreadable codons in essential positions. Second, genetic-code-sensitive gene expression of kill switches or reporters will counterselect for the tendency to reincorporate the wild-type amino acid in TCR or other unassigned codons. Third, following a previously defined strategy^19^, we propose to make the recoded cell dependent on the exogenous supply of a synthetic non-standard amino acid (nsAA) like biphenylalanine (bipA). This strategy involves protein engineering to make an essential protein dependent on bipA. The larger the number of essential proteins depending on bipA, the lower the escape rate. Fourth, the remaining free codons could be reassigned to new non-standard amino acids to expand the biochemical applications of cells. In this figure, tRNA^Ser^ refers to serine tRNAs with YGA anticodon that recognize TCR codons.

Our strategy involving genetic-code-sensitive kill switches and reporters expands biocontainment and applications of prior efforts to reassign codons to natural amino acids^20,21^. Specifically, we prevent recoded cells from escaping a genetic-code-based bioisolation via tRNA modifications or mutations (base changes) linking serine and TCR codons, while we simultaneously allow non-standard amino acid incorporation. Several genetic-code-sensitive kill switches spread throughout the genome could be combined with alternative technologies to further improve the long-term stability of a genetic-code-based cellular bioisolation. An alternative approach to increase the life span of a kill switch could entangle an essential gene with a genetic-code-sensitive antibacterial, building on previous suggestions overlapping genes and toxins^52,53^. Another option to reduce likelihood of escape involves the proteome-wide incorporation of new standard or non-standard amino acids into the reassigned codons. The dependency of many essential enzymes on the non-standard amino acid would stabilize the reassigned codon in the corresponding essential genes (Supplementary Fig. 16b). This aspect is critical to prevent the return of a wild-type link between an unassigned sense codon and its natural amino acid. Feasibility of this approach has been supported by previous work showing that the incorporation of non-standard amino acids in amber stop codons within a handful of essential enzymes^19,54^ rendered the lowest escape rate to-date in a living organism, *i.e.*, 10^-12^ cfu/mL.

Engineering tens or hundreds of essential proteins to depend on a non-standard amino acid in a recoded cell remains a promising, stabilizing biocontainment goal. To design fit proteins containing non-standard amino acids, we need the guidance of data and new machine learning models that expand the limitations of those currently available^55,56^. We expect that reassigning sense codons naturally linked to distinct amino acids (as ongoing in our upcoming *E. coli* strain with a 57-codon genome^3^ or Ec_Syn57Δ6, Supplementary table 1) will enable both tight bioisolation and incorporation of multiple non-standard amino acids. Ultimately, living technologies need multiple layers of biocontainment to ensure safe use. These layers could involve genomic recoding, knockout of non-essential promiscuous aminoacyl-tRNA synthetases, genetic-code-sensitive kill switches and reporters, and a genome-wide codon reassignment to incorporate non-standard amino acids into essential proteins (Fig. 4). Having these layers will allow for controlled manipulation of the genome so to direct cell function precisely and responsibly.

As shown in this work, genetic-code-engineered cells can be used to study the crosstalk, evolvability, and biosafety of new genetic codes, in addition to offering a promising approach to living technologies. “Sense” codon recoding allows the incorporation of multiple non-standard amino acids, which promises to expand engineered protein applications significantly. Nevertheless, recoded sense codons without a cognate tRNA might be less likely to inhibit translation than a recoded amber stop codon^1,57^. Innate cellular mechanisms like tmRNAs or promiscuous aminoacyl-tRNA synthetases act during translation to resolve problems like ribosome stalling and ensure cellular survival^57^. While changes in tRNA expression^58^, sequence^59–61^, and modifications^38,62^ seem to be widespread in nature as a viral defense mechanism or to change suppression efficiencies, we show for the first time the essential role that non-cognate tRNAs play in the bioisolation of a genetic code engineered organism. Our work has broad implications for the understanding of the genetic code, the future design, biosafety of synthetic genomes, and the incorporation of non-standard amino acids.

## Methods

### Bacterial liquid media

We prepared both liquid Luria Broth (LB) and liquid Super Optimal Broth (SOB) medium from LB Medium Lennox Capsules (76019-956, MP Biomedicals) and from Difco SOB Medium powder (244310, BD Diagnostics), respectively. We added and mixed 10 capsules of LB Medium Lennox Capsules or 14 g of SOB Medium powder, respectively, in 500 mL of dH_2_O. We autoclaved all bottles with media at 121°C for 45 minutes in a liquid cycle and stored them at room temperature.

### Bacterial LB agar medium preparation

We mixed 4 L of dH_2_O with 80 g of Difco Bacto Agar (214030, Becxton Dickenson), 40 g of Bacto Tryptone Pencreatic Digest of Casein (211699, Gibico/ LifeTechnologies Co.), 20 g of Bacto Yeast Extract (212720, Gibico/ LifeTechnologies Co.), 20 g of Sodium Chloride NaCl (0241-10kg, VWR Life Sciences). We autoclaved all bottles with agar media at 121°C for 45 minutes in a liquid cycle. After autoclaving, we let the LB agar cool down to approximately 30°C. We added the required amounts of reagents from stock solutions (*e.g.*, antibiotics, glucose, arabinose, bipA) to achieve the target concentrations. We plated 20-25 mL of LB agar onto each culture plate. The poured LB agar medium cooled down under a laminar flow hood (HeraguardECO, Thermo Fisher Scientific) for 45 minutes. We packed and stored the plates at 4°C until use.

### Antibiotics and concentrations

We worked with the following antibiotics and final concentrations: Chloramphenicol (C0316, Teknova) at 25 µg/mL, Kanamycin (K2199, Teknova) at 50 µg/mL, Carbenicillin (E0096, Teknova) at 100 µg/mL, Spectinomycin (S9525, Teknova) at 75 µg/mL, and Gentamicin (15750060, Thermo Fisher Scientific, Life Technologies) at 15 µg/mL. We defined each antibiotic with a letter code: A for chloramphenicol, B for kanamycin, C for carbenicillin, D for spectinomycin, and E for gentamicin. We stored all antibiotic stock solutions at -20°C, except for gentamicin, which was stored at 4°C.

### L-4,4’-Biphenylalanine (bipA) and concentrations

We worked with bipA at a final standard concentration of 100 µM. We prepared a 100 mM stock solution of bipA by dissolving 482 mg of bipA powder (155760-02-4, Astatech) in 18 mL of milliQ H_2_O, adding 600 µL of 10M NaOH, and bringing the final volume up to 20 mL with milliQ H_2_O. We filter sterilized the bipA solution with a 0.22 µm syringe filter. We stored bipA stock solutions in 1 mL aliquots at -20°C.

### Electrocompetent cell preparation of Ec_Syn61Δ3

We prepared big batches (∼100 final aliquots) of competent cells with the same concentration of Ec_Syn61Δ3 to allow obtaining pairs of replicates for all plasmid designs with each competent cell batch. We added 500 mL of Super Optimal Broth (SOB) without antibiotics in a 2 L flask in aseptic conditions under a laminar flow hood (HeraguardECO, Thermo Fisher Scientific). We worked with a starting optical density (OD_600_) of 0.003 in 500 mL of SOB from the overnight cultures. We grew cells in a shaking incubator at 37°C and 250 rpm in aerobic conditions until the OD_600_ reached between 0.1-0.4. When ready, we placed the flask immediately on ice for a fast cool down for 15 minutes. The next steps were done without pauses and trying to prevent heating of the competent cells. We transferred the culture to pre-cooled 50 mL falcon tubes. We centrifuged for 10 minutes at 4500×g and 4°C. We discarded the supernatant and resuspended the pellets with 1 mL of pre-cooled sterile dH_2_O with 10% glycerol, transferring the content to a pre-cooled Eppendorf tube. Next, we centrifuged for 2 minutes at 5000×g and 4°C. We repeated the resuspension and centrifugation steps four times. In the last round we resuspended each pellet in approximately 350 µL of pre-cooled dH_2_O with 10% glycerol (volume estimated based on size of pellet). We collected the whole volume with competent cells in a new precooled 50 mL falcon tube to assure all subsequent aliquots had the same concentration of cells. We transferred 40 µL of cells to pre-cooled Eppendorf tubes and stored them immediately at -80°C.

### Plasmid electroporation

We added 200 ng in 1 µL of plasmid suspended in IDTE (76286-184, G-Biosciences) to the 40 µL of competent cells. We pipetted gently up and down 3-5 times for careful mixing and transferred the mix to a pre-cooled electroporation cuvette (1652089, Bio Rad Laboratories). We applied the standard electroporation program for *E. coli*: 1.8 kV, 200 Ω, 25uF, 1 mm in a Bio-Rad Gene Pulser Xcell electroporation machine (1652660, Bio-Rad). We collected the electroporated culture with 1 mL of SOB pre-warmed at 37°C and transferred it to a 13 mL culture tube with vented cap. The culture grew overnight in a rotator at 37°C.

### Plating and escapee observation

We added sterile ColiRollers plating beads to pre-warmed Luria Broth (LB) agar plates containing or not antibiotics. For identification of escapees, we plated 800 µL of overnight culture without dilution, distributing 100 µL of culture per plate. For the electroporation of plasmids with a compatible genetic code to the host cell or for escapees with a high escape rate, we prepared dilutions 1:1, 1:10, 1:10^2^, 1:10^3^, 1:10^4^, 1:10^5^, 1:10^6^ from the overnight culture in liquid SOB. We plated 100 µL of these dilutions onto plates. We moved the plates gently to spread the culture with the beads. The plates dried under a laminar flow hood (HeraguardECO, Thermo Fisher Scientific) for approximately 10 minutes. We discarded the beads and placed the plates in an incubator at 37°C. The plates remained in the incubator until visible colonies were formed, which took 2-3 days for Ec_Syn61Δ3 escapees in most scenarios. When no escapees were seen in 3 days, plates were left longer in the incubator for up to 10 days. Colonies were inoculated from plates incubated for up to 5 days. We either washed these plates with PBS for bulk analysis of escapees or inoculated colonies from these plates for individual analysis of escapees, as described next. Plates with colonies were also imaged for colony quantification, as described next.

### Bacterial escape rate measurement

We quantified the number of cells in each electrocompetent cell batch by plating dilutions of overnight cultures after electroporation onto LB agar. We plated 20 µL of dilutions 1:10^3^ and 1:10^5^ from the overnight culture onto LB for colony quantification. Although the initial number of competent cells before electroporation was in the order of 10^9^ cfu/mL, the cfu/mL onto LB agar plates by a plasmid-less overnight culture of Ec_Syn61Δ3 that underwent electroporation was measured to be in the order of 10^7^ cfu/mL for all competent cell batches. All reported measurements of colonies and escape rates in cfu/mL were normalized to a number of competent cells of 2.1x10^7^ cfu/mL. We did this to be able to compare escapees relative to the same number of competent cells across replicates (Fig. 1d). It is important to note, that with the approach followed, measurements of escape rate are affected by the plasmid uptake efficiency during electroporation, and hence it is expected that the actual escape rates are higher than the values reported.

To eliminate the component of the electroporation efficiency in the escape rate measurement, we also measured the colonies formed (cfu/mL) by Ec_Syn61Δ3 with the recoded plasmid pUC57-dTCR (Fig. 1c), which defined the expected number of colonies with a genetic code compatible plasmid. We defined a relative escape rate [(cfu/mL)/ (cfu/mL)] as the number of colonies formed by Ec_Syn61Δ3 with the wild-type plasmid pUC57 relative to those of parallel experiments with pUC57-dTCR upon selection onto carbenicillin-containing agar plates.

All plates with colonies were quantified by hand. For conditions rendering hundreds of colonies or above, we divided the plate into eight sections of the same area and counted the number of colonies in one section (*C*_1/8*P*_). The number of colonies per mL was next calculated as

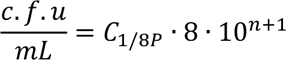

where *n* is the dilution of the plate in which the colonies were counted. We report the logarithmic values in base 10 of the cfu/mL. We also calculated a relative escape rate of Ec_Syn61Δ3 with wild-type pUC57 compared to the recoded plasmid pUC57dTCR. For this purpose, we divided the colonies per mL formed with the wild-type pUC57 and the recoded pUC57dTCR. We validated some measurements with a colony counting pipeline developed within the software CellProfiler^63^.

### Quantification of escape rate by plasmid-less Ec_Syn61Δ3

We quantified colonies formed by plasmid-less Ec_Syn61Δ3 with two tests. In the first test, Ec_Syn61Δ3 underwent electroporation (without plasmid), overnight growth, plating, and selection onto agar plates containing standard antibiotic concentrations, following the same approach as experiments in which escapees were observed (previous section). We plated 800 µL of overnight culture, distributing 100 µL per plate. We also plated 20 µL of dilutions 1:10^3^ and 1:10^5^ from the overnight culture onto LB for colony quantification. In the second test, we grew Ec_Syn61Δ3 in liquid LB without antibiotics overnight. We normalized the cultures for an OD_600_ = 1 and plated 1 mL of the culture in antibiotic containing agar plates, distributing 100 µL per plate. Antibiotic-containing agar plates were incubated for 10 days at 37°C, and LB plates were incubated for 3 days at 37°C.

### Plasmid extraction and sequence analysis

We spun down 1-3 mL of culture for plasmid extraction, discarded the supernatant and stored the cell pellet at -20°C. We used the Qiagen miniprep kit protocol for plasmid extraction. In the last step, we eluted the plasmid with 50 µL of IDTE pre-warmed at 45°C. We sent plasmids for long read nanopore sequencing. Raw reads were mapped to the reference map with the plasmid design (Supplementary Datasets 1 and 2) for sequence validation. We used the Geneious mapper and Bowtie2 with a medium sensitivity within Geneious Prime (versions 2022-2024) for mapping of raw sequences to all plasmid maps. We also sent 100 µL of overnight cultures or cell pellets resuspended in 10 µL of dH_2_O for zero-prep whole plasmid sequencing. Raw plasmid sequences are provided under the NCBI bioproject ID defined in the Data Availability section.

### Bacterial genome sequencing and sequence analysis

We spun down 1 mL of culture for genome sequencing, discarded the supernatant and stored the cell pellet at -20°C. We sent cell pellets in dry ice for bacterial genomic DNA extraction and Illumina whole genome sequencing. We chose a sequencing depth or coverage of 200 Mbp or approximately 1.33M Reads. For specific samples, we increased the sequencing depth or coverage to 5 Gbp or approximately 33.3M Reads. Raw reads were mapped to two genomes (Supplementary Dataset 3): (1) to an *E. coli* genome with *serT* and *serU* genes, *i.e.*, MDS42, for sequence validation and confirmation that *serT* and *serU* genes were not present in the strains; (2) to the Ec_Syn61Δ3 for sequence validation and identification of single nucleotide mutations. We used the Geneious mapper and Bowtie2 with a medium sensitivity within Geneious Prime (versions 2022-2024) for mapping of raw sequences to both genomes. We identified the percentage of reads mapped, which seems to inversely correlate with an increase in duplicated genomic regions. We looked for an increase in the gene copy number based on the genome coverage. Specifically, we identified genomic loci with an increase in the number of reads mapped to it above 2 of the standard deviation from the mean coverage. We also looked for single nucleotide polymorphisms (SNPs) in the genome of MDS42 with mapped sequencing data from escapees. We defined as SNPs those genomic mutations that: (1) did not involve recoding substitutions TCA->AGT or TCG->AGC; (2) were present in escapees and not in MDS42 with mapped sequencing data from plasmid-less Ec_Syn61Δ3; (3) were present at an 80% or higher frequency. We identified SNPs common to Ec_Syn61Δ3 escapees containing the same plasmid and combined all unique SNPs (Supplementary table 4). We visually analyzed location of SNPs in the genome maps. The MDS42 and Ec_Syn61 Δ3 genome maps used are provided in Supplementary Dataset 3. All raw genome sequences are provided under the NCBI bioproject ID defined in the Data Availability section.

### Sample acquisition for tRNA-seq, proteomics, plasmid and genome sequencing, *in vivo* **reporting of tRNA mutations**

We took three types of samples of Ec_Syn61Δ3 escapees. Escapees were first observed on LB agar plates with the corresponding antibiotics after electroporation with plasmids containing antibiotic resistance genes.

*Option 1*. We washed the plate with colonies of escapees with 2 mL of PBS. We transferred the mix of PBS with escapees to an Eppendorf tube and centrifuged it at 6000×g for 2 minutes. We discarded the supernatant, washed the pellet twice more with 1 mL of PBS, and centrifuged the samples again at 6000×g for 2 minutes. We discarded the supernatant and stored the cell pellets at -20°C for further downstream analysis.

*Option 2*. We added 5 mL of LB with the corresponding standard antibiotic concentration in 13 mL culture tubes. We inoculated colonies of Ec_Syn61Δ3 escapees from LB antibiotic-containing agar plates separately into each culture tube. These cultures grew for one to three days in a rotator at 37°C. We used 1 mL of escapees’ cultures for stocks and transferred 1 mL of culture to Eppendorf tubes for downstream analysis: genome sequencing, plasmid sequencing, and his-tag pull down for proteomics. We used the rest of escapees’ cultures for Option 3 and for dynamic fluorescence/growth tracking with our genetic-code-sensitive reporter. We centrifuged the cultures in Eppendorf tubes at 6000×g for 1 minute. Cell pellets for proteomics were washed three times with 1 mL of PBS. We discarded the supernatant and stored the pellets at -20°C for further processing depending on the downstream analysis.

*Option 3*. We added 50 mL of LB with the corresponding standard antibiotic concentration and L-arabinose for a final 0.2%-v/v concentration in 250 mL flasks. We transferred 50 µL of escapee’s culture (from Option 2) to each flask. This resulted in an approximate starting optical density of OD_600_ 0.001. We grew these cultures in a shaking incubator at 37°C and 250 rpm in aerobic conditions until exponential phase with OD_600_ between 0.3 and 0.7. We took samples of these cultures for genome sequencing (1 mL), plasmid sequencing (2 mL), tRNA-seq (15 mL), and proteomics (5 mL). We centrifuged 50 mL falcon tubes for 5 minutes at 4500×g and Eppendorf tubes for 1 minute at 6000×g. Cell pellets for proteomics were washed three times with 1 mL of PBS. We discarded the supernatant and stored the cell pellets at - 20°C.

### Dynamic growth and fluorescence assays

We prepared a 96-well plate with clear flat and black bottom polystyrene (3603, Corning Life Sciences) with 200 µL of LB with the corresponding standard antibiotic concentration and when required 1%-v/v of D-glucose, or 0.2%-v/v of L-arabinose, or 100 µM of bipA. D-glucose represses expression of our genetic-code-sensitive genes, while L-arabinose induces expression of our genetic-code-sensitive genes. We transferred 2 µL of overnight culture (*e.g.*, escapees’ culture from Option 2 above or a culture of Ec_Syn61 Δ3 with genetic-code-sensitive kill switch normalized to OD_600_ = 1) into each well for a 1:100 dilution. We set up the plate with a lid in a microplate reader (Synergy H1 BioTek). We took measurements every 10 minutes of absorbance at wavelength 600 nm to track growth through optical density (OD_600_) and of excitation/emission at 485/528 nm with gain 100 or 50 to detect fluorescence. This program ran for 48 or 72 hours at 37°C and with linear shaking. Data from these assays is provided in Supplementary table 7. Growth rates were estimated by identifying the best linear fit of maximum positive slope in-between intervals of 3.3 hours. The linear fit and R-square were plotted and manually validated.

### Cell Specific Sample Preparation for untargeted proteomics

Frozen cell pellets were dissolved for 15 minutes in 5 % SDS buffer at room temperature to lyse cells. The solution was run on a micro S-trap column (ProtiFi, NY) according to the manufacturer protocol (Protocols – ProtiFi)^64^.

### His-tag pull down for targeted proteomics

Frozen cell pellets were dissolved in 2 mL of BugBuster Protein Extraction Reagent (70584-3, MilliporeSigma) and incubated at room temperature for 5 minutes. The lysed cell mixture was spun down at 8,000 rpm for 10 minutes. The supernatant was transferred to a new tube and mixed with 2 mL of His-Binding/Wash Buffer (786-542, G-Biosciences) and 5 μL of HisTag Dynabeads (10104D, Thermo Fischer Scientific). This mixture incubated at room temperature for 5 minutes. The beads were separated on a magnetic rack and the liquid was pipetted out and discarded. The beads were washed with 300 μL of His-Binding/Wash Buffer (786-542, G-Biosciences) and phosphate-buffered saline (1X PBS) three times. After the final wash, all liquid was removed and the bead pellets containing the protein samples was frozen down at −80°C.

### TMT Labeling

Samples, now peptides, were labeled using 10uL of TMTpro Mass Tag Labels (ThermoScientific). After labeling, samples were then placed on Eppendorf ThermoMixer C for 45 minutes and shaken/mixed for 45 minutes; this ensures labels are covalently bonded to peptides. Labeling reaction was quenched for 10 minutes using 1 µL of 5% hydroxyalamine. After quenching, samples were pooled into one 2 mL Eppendorf tube and then dried down using Eppendorf Vacufuge plus.

### Hi pH separation of TMT labeled material

Dried samples were resuspended in 20 µL of 0.1% TFA in ultrapure HPLC (High Pressure Liquid Chromatography) grade water and vortexed to ensure full solubility. Before submission to LC-MS/MS experiment sample was separated on Hi-pH column (ThermoScientific, CA) according to vendor instructions to 20 fractions. Desalted samples were then transferred to HPLC (High Pressure Liquid Chromatography) vials and were dried down using Eppendorf Vacufuge plus. Dried and desalted samples were resuspended in 6 µL of 0.1% Formic Acid in ultrapure HPLC (High Pressure Liquid Chromatography) grade water and were then injected on LC-MS/MS (Liquid Chromatography tandem Mass Spectrometry).

### Mass spectrometry analysis

After separation, each fraction was submitted for single LC-MS/MS experiment that was performed on a Q Exactive Orbitrap (Thermo Scientific, CA, USA) equipped Evosep (Odense, Denmark) nanoHPLC pump. Peptides were separated onto a 150 µm x 8 cm PepSep C18 analytical column (Bruker, MA). Separation was achieved through applying a gradient from 5– 25% ACN in 0.1% formic acid over 45 minutes at 300 nL/min. Electrospray ionization was enabled through applying a voltage of 2 kV using a PepSep electrode junction at the end of the analytical column and sprayed from stainless still PepSep emitter SS 30 µm (Bruker, MA). The Q Exactive Orbitrap was operated in data-dependent mode for the mass spectrometry methods. The mass spectrometry survey scan was performed in the Orbitrap in the range of 450 –900 m/z at a resolution of 1.2 × 10^5^, followed by the selection of the ten most intense ions (TOP10) ions were subjected to HCD MS2 event in Orbitrap part of the instrument. The fragment ion isolation width was set to 0.8 m/z, AGC was set to 50,000, the maximum ion time was 150 ms, normalized collision energy was set to 34V and an activation time of 1 ms for each HCD MS2 scan.

### Mass spectrometry data analysis

Raw data were submitted for analysis in Proteome Discoverer 3.1.0.638 (Thermo Scientific) software with Chimerys 2.0. Assignment of MS/MS spectra was performed using the Sequest HT algorithm and Chimerys (MSAID, Germany) by searching the data against a protein sequence database including all entries from the *E. coli* K12 database and other manually curated databases with all possible new protein insertions as well as known contaminants such as human keratins and common lab contaminants. Sequest HT searches were performed using a 10 ppm precursor ion tolerance and requiring each peptides N-/C termini to adhere with Trypsin protease specificity, while allowing up to two missed cleavages. 18-plex TMT tags on peptide N termini and lysine residues (+304.207146 Da) was set as static modifications and Carbamidomethyl on cysteine amino acids (+57.021464 Da) while methionine oxidation (+15.99492 Da) was set as variable modification. Specific amino acid modification to Phenyl alanine amino acid was made as an addition of C6H4 (+ 76.09617 Da) to represent bipA synthetic amino acid. A MS2 spectra assignment false discovery rate (FDR) of 1% on protein level was achieved by applying the target-decoy database search. Filtering was performed using a Percolator (64bit version^65^). For quantification, a 0.02 m/z window centered on the theoretical m/z value of each of the six reporter ions and the intensity of the signal closest to the theoretical m/z value was recorded. Reporter ion intensities were exported in result file of Proteome Discoverer 3.1 search engine as an excel tables. The total signal intensity across all peptides quantified was summed for each TMT channel, and all intensity values were adjusted to account for potentially uneven TMT labeling and/or sample handling variance for each labeled channel.

### Amplification of *serT* and *serU* genomic regions and Sanger sequencing

For each PCR reaction we mixed 11.5 μL of 2x Platinum II Hot-Start Green PCR master mix (14001012, Invitrogen), 0.06 μL of forward primer (100 μM), 0.06 μL of reverse primer (100 μM), 12.5 μL of dH_2_O, and 1 μL of sample (liquid culture or colony disolved in 5 μL of LB and 5 μL of dH_2_O). Our PCR protocol involved one cycle with 94°C (five minutes), 28 cycles with 94°C (20 seconds), 61°C (20 seconds), and 68°C (2 minutes), and a last cycle at 68°C (4 minutes). To evaluate whether there is any contamination of cells containing *serT* and/or *serU*, we used the following primers which should not give a band in Ec_Syn61Δ3.

F_serT_int: TCGAACTCTGGAACCCTTTCG (paired with R_serT_ext)

R_serT_ext: CTTATGGTCGGCAAGCGAC

F_serU_int: GGAGAGAGGGGGATTTGAACC (paired with R_serU_ext)

R_serU_ext: CGGGGCGAATATTACTTAGCG

To further confirm the sequence of the genomic region surrounding the loci where *serT* and *serU*, we used the following external primers:

R_serT_ext: CTTATGGTCGGCAAGCGAC (pair with F_serT_ext)

F_serT_ext: GCACCCAGAAGTTTTACCCATC

R_serU_ext: CGGGGCGAATATTACTTAGCG (pair with F_serU_ext)

F_serU_ext: CTTTGCGGCAGTTTCATAGGA

Amplicons from external primers were sent for Sanger sequencing. Raw reads were mapped to an *E. coli* genome with *serT* and *serU* genes, *i.e.*, MDS42, for sequence validation and confirmation that *serT* and *serU* were not present in the strains. We used the Geneious mapper with a medium sensitivity within Geneious Prime (versions 2022-2024) for mapping of raw sequences to MDS42 genomic regions surrounding *serT* and *serU*. Gel images are provided in Supplementary Fig. 5.

### Design of a bipARS/tRNA pair that incorporates bipA into TCG codons

We mutated the anticodon of tRNA^bipA^ to be CGA and thereby be able to recognize TCG codons for bipA incorporation as tRNA^bipA^_CGA_. We tested incorporation of bipA with the new tRNA^bipA^_CGA_ and a library of twenty-one bipARS variants. We used the wild-type bipARS sequence and also mutagenized bipARS at positions 286 (D286R, based on previous literature^66^) and 288 (K288N, based on the crystal structure of MjTyrRS^66,67^ and the potential interaction of this amino acid position with the second nucleotide of the tRNA^bipA^ anticodon). These mutant genes did not contain TCR codons. The tRNA^bipA^_CGA_ and bipARS variants (the latter under an arabinose inducible promoter) were cloned in the pEVOL backbone (same as, *e.g.,* Addgene plasmid #73546). Sequences are provided in Supplementary table 9 and Dataset 2.

### Sample preparation of bipARS library for Illumina sequencing

The DNA from the cell pellets was extracted via Midiprep and diluted to 0.5 ng/µL. A PCR reaction with 10.35 μL of the diluted Midiprep, 1.15 μL of the UMI primer (primer 1) and 11.5 μL of 2x Platinum II Hot-Start Green PCR master mix (14001012, Invitrogen) was set up. One cycle with 94°C (two minutes), 61°C (15 minutes), and 68°C (15 minutes) was run to attach the UMI primer. Subsequently, we added 1 μL of Primer 2 and Prime 3 each to the reaction, together with 13.5 μL of 2x Platinum PCR master Mix and 11.5 μL of H2O and mixed by pipetting. The resulting 50 μL reactions were subjected to 94°C (two minutes) and 30 cycles of 94°C (15 seconds), 61°C (15 seconds), and 68°C (30 seconds), followed by 68°C (4 minutes). The PCR products were purified with AMPure XP beads, following the standard protocol. The resulting DNA concentration was determined by Qubit and diluted to 20 ng/μL. <u>Primers:</u>

Primer 1: ACACTCTTTCCCTACACGACGCTCTTCCGATCTNNNNNggatctgaccgtcaactccta

Primer 2: GACTGGAGTTCAGACGTGTGCTCTTCCGATCTNNNNNccttatccggcctacaaaagc

Primer 3: ACACTCTTTCCCTACACGAC

### Processing Illumina sequencing reads

The obtained Illumina reads from Azenta were processed with FLASH-1.2.4 to merge read 1 and read 2. The output was passed into seqkit to reverse complement and filter for mean quality score of 20. Using a custom script, the relevant region with the library was extracted and translated into amino acids. Amino acid frequencies were calculated and averaged across three replicates. By dividing the amino acid frequency with Arabinose (induction of bipARS expression) by the amino acid frequency in glucose (inhibits bipARS expression), we obtained enrichment scores. Those enrichments were converted into log2-enrichments. Comparing the log2-enrichment score of samples with and without bipA added, we obtained a fractional difference score for the effect of adding bipA and used that to rank variants.

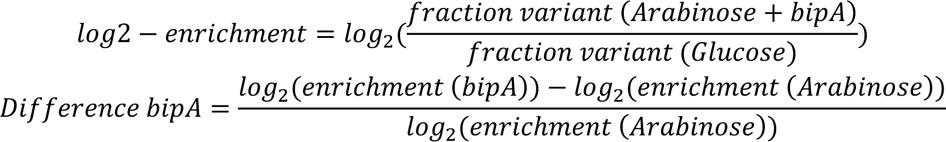

### Database for proteomics analysis of bipA incorporation

To query proteins with incorporation of amino acids at TCG codons, we split each sequence with multiple TCG codons into multiple fragments. Splitting those fragments allowed, *e.g.*, sampling all pairwise combinations between these two or more TCA sites without having a large database. We only separated two TCG positions, if at least three amino acids were separating the two positions. Otherwise, the two positions remained in one fragment. For the separation we made overlapping ends, meaning that after and before each incorporation site was the end of the fragment leading to overlap regions to the next fragment. We created all combinations of possible incorporated amino acids at these sites in all fragments. This fragment database was added to a general database of *E. coli* genes to match fragments identified in mass spectrometry.

### Database for proteomics analysis of serine incorporation

We generated sequences of all target proteins as follows. For protein sequences whose genes contain three or less TCR codons, we generated protein sequences with all combinations of amino acid mutations in those TCR positions. We mutated those positions to all the 20 standard amino acids. For protein sequences whose genes contain more than three TCR codons, we could not follow the previous approach because the database could not be handled by the proteomics analysis software. We generated protein sequences with all possible amino acid mutations in each TCR codon position separately. If two TCR codons were separated by less than 10 amino acids, we generated protein sequences with all combinations of amino acid mutations in these positions. We mutated those positions to all the 20 standard amino acids.

### Design of genetic-code-sensitive gene expression based on protein language models

We used a protein language model called Evolutionary Scale Model (ESM-1b)^32^ from Facebook Research available at https://huggingface.co/facebook/esm1b_t33_650M_UR50S and https://github.com/facebookresearch/esm. Protein language models predict the probability that an amino acid like serine is present in a position *n* (*P*(*Ser*_*n*_)) to render a functional protein. We ranked these probabilities to identify the top ranked positions and the lowest ranked positions. We hypothesized that Ec_Syn61Δ3 can initially escape through mistranslation of antibiotic resistance genes, which depends on the number (*N*) and positions (*n*) of TCR codons, as we observed (Supplementary Fig. 3). We defined TCR codons in top or lowest ranked serine positions for our genetic-code-sensitive gene designs. These aimed to render the highest and lowest probability of expression in Ec_Syn61Δ3. We quantified the probability of expression *PE* or theoretical biocontainment for a genetic-code-sensitive gene design as follows. A genetic-code-sensitive gene design contains *N* TCR codons for *N* serine positions. We assumed that the misincorporation of amino acids at each of the *N* TCR-coded serine positions within a protein is due to independent events. Hence, independent misincorporation probabilities (1 − *P*(*Ser_n_*)) could be multiplied to estimate an overall probability of mistranslation and escape (*PE*).

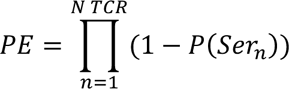

### Design of antibiotic resistance genes

We developed three types of designs for each gene conferring resistance to chloramphenicol, kanamycin, carbenicillin, spectinomycin, and gentamicin. All these designs are his-tagged at the C-terminus, except for the kanR design which needed to be tagged at the N-terminus due protein fitness problems. Each design was called as pX_design type_, where *p* denotes plasmid, *X* is the letter code used for each antibiotic, and *design type* refers to the procedure used to design the gene. Designs tagged as “wild-type” contain TCA codons in the same positions where a wild-type gene design contains TCR codons. Designs tagged as “max” contain TCA codons in the serine positions with highest probability for serine based on the protein language model ESM-1b. Designs tagged as “min” contain TCA codons in the serine positions with lowest probability for serine based on the protein language model ESM-1b. We kept constitutive promoters and ribosome binding sites from available plasmids with wild-type antibiotic resistance genes like pUC57 (Supplementary table 2, Supplementary Dataset 1).

### Design of genetic-code-sensitive reporter

We designed a genetic-code-sensitive GFP reporter gene that contains five TCA codons in the serine positions with highest probability for serine based on the protein language model ESM-1b. This gene is expressed under an arabinose-inducible promoter and is his-tagged at the C-terminus. It was cloned in the plasmids containing our designs of antibiotic resistance genes (Supplementary table 8).

### Design of genetic-code-sensitive kill switches

We selected toxins from the literature^22,43,45^ with known toxicity against *E. coli*. We selected 3-14 wild-type serine positions in which serine is the most likely amino acid to render a functional protein based on the protein language model ESM-1b^32^ (Supplementary table 8). We defined those codons as TCG or TCA in the gene sequence. We designed a strong ribosome binding site using deNovoDNA^68^ (https://www.denovodna.com/software/design_rbs_calculator) and added a strong constitutive promoter upstream for expression. The designs were synthesized and cloned into the recoded backbone vector with pUC or pA15 origins of replication.

### Toxin database for analysis and visualization

We downloaded the toxins from the TADB3.0 database^42^ (access February 2024) and extracted all amino acid sequences from the subset with experimental validations and the subset from in-silico predictions. After removing duplicate toxins based on the amino acid sequence, this dataset resulted in a total of 2120 toxins (2058 from the TADB in-silico subset, 62 from the experimental validation subset). We added to this data 11 hand curated toxin sequences from the literature^22,43,45^.

### Toxin 2D visualization

The vector representation of each toxin was created by using a protein language model (ESM-2, https://github.com/facebookresearch/esm)^69^. For each toxin, we computed the mean over the hidden representations of the last model’s layer for each amino acid in the toxin amino acid sequence, resulting in vectors of dimensions 1x2058 for each toxin. To represent the toxin data in 2D, we used the t-distributed stochastic neighbor embedding (t-SNE) method^70^ to project the extracted ESM-2 toxin representation in 2D using the function TSNE from the Python library scikit-learn (https://scikit-learn.org/stable/) with the default parameters.

### Clustering toxin sequences

To cluster the toxins in groups, we used the Density-based spatial clustering of applications with noise (DBSCAN)^71^. We used the implementation from the Python library scikit-learn (https://scikit-learn.org/stable/) with the following parameters:

The maximum distance between two samples for one to be considered as in the neighborhood of the other (eps): 1.6

The number of samples (or total weight) in a neighborhood for a point to be considered as a core point (min_samples): 5

This resulted in 30 clusters (colored dots in Supplementary Fig. 13) and 45 toxin sequences that could not be assigned to a cluster (not shown in Supplementary Fig. 13).

### Functional evaluation of kill switches

We tested the functionality of our kill switches in four ways. First, we tested that our kill switches were not functional without a tRNA^Ser^ decoding TCR codons. We worked with a strain of Ec_Syn61Δ3 containing the plasmid with the kill switch. We evaluated the growth curves in a plate reader between Ec_Syn61Δ3 containing or not the plasmid with the kill switch. Second, we tested that our kill switches were not functional in the presence of an aminoacyl-tRNA synthetase and tRNA pair able to incorporate bipA into TCG codons (bipARS/tRNA^bipA^_CGA_). The aminoacyl-tRNA synthetase is expressed with an arabinose inducible promoter. We worked with a strain of Ec_Syn61Δ3 containing both the plasmid with the kill switch and the plasmid with the bipARS/tRNA^bipA^_CGA_ system. And we compared growth of the strain in a plate reader between a medium that included 1% D-glucose or 0.2% L-arabinose. Third, we tested that the kill switch was expressed in the presence of a tRNA^Ser^ serT and serU. We placed the tRNA^Ser^ under an arabinose inducible promoter. We worked with a strain of Ec_Syn61Δ3 containing both the plasmid with the kill switch and the plasmid with the inducible tRNA^Ser^ expression system. And we compared growth of the strain in liquid medium and onto solid medium that included 1% D-glucose or 0.2% L-arabinose. Fourth, we evaluated the escape rate of the strain Ec_Syn61Δ3 with a kill switch in the presence of antibiotic resistance genes containing TCR codons. We electroporated a plasmid containing a Kanamycin resistance (KanR) gene with four TCR codons and a plasmid with the genetic-code-sensitive kill switch. We selected onto an LB agar plate containing carbenicillin and kanamycin antibiotics. Plates were incubated for up to 10 days at 37°C.

### RNA isolation

Cell pellets were suspended in 1mL TRIzol™ (Thermo Fisher) and frozen at -80°. 1/10 volume 1-bromo-3-chloropropane were added and the samples were vortexed and centrifuged at 15,000×g for 15 minutes at 4°C. The aqueous phase was transferred to a new tube containing 400 μL of 70% ethanol. Short RNAs were then isolated using a modified RNeasy MinElute Cleanup Kit protocol (Qiagen) to size select for RNAs <200 nt in length. Samples were centrifuged through MinElute spin column at 12,000×g for 5 minutes at room temperature. The flow-through was added to 450 μL of 100% ethanol and centrifuged in a new MinElute spin column at 12,000×g for 1 minute at room temperature. The column was washed 3 times with 80% ethanol in 50 mM sodium acetate and dried with open caps at 12,000×g for 5 minutes. Samples were eluted in 50 mM sodium acetate and 1 mM EDTA.

### tRNA library preparation and sequencing

tRNAs were ligated to a 3’ pre-adenylated adaptor oligo using New Englad Biolabs T4 RNA Ligase 2, truncated KQ kit to ligate adenylated 5’ end of a DNA oligo to the 3’ OH on RNA samples. Reaction was carried out according to manufacturer’s protocol for 2 hours at 25°C and quenched with 0.5 M EDTA, and subsequently size-selected with SPRIselect. Primer-dependent reverse transcription reaction was carried out using Maxima H Minus Reverse Transcriptase. 50 ng of 3’-adaptor ligated samples were incubated with 6 pM RT primer and 20 mM dNTP mixture at 75°C for 5 minutes to denature primer and added to Maxima kit components according to manufacturer’s protocol. After incubation, alkaline hydrolysis was carried out through the addition of NaOH and incubation at 95°C for 3 minutes, and solution was neutralized with HCl. Samples were purified using SPRIselect. 5’ adaptor was ligated using New Englad Biolabs Thermostable 5’ App DNA/RNA Ligase according to manufacturer protocol, and adaptor-ligated samples were size selected with SPRIselect. Barcodes were added to 3’ and 5’ ends of cDNA through single-step PCR with KAPA HiFi HotStart ReadyMix using primers including the Illumina P5 and P7 regions and barcodes adapted from Illumina TruSeq. Libraries were purified using SPRIselect and quantified using QuBit. Libraries were prepared for sequencing using Illumina MiSeq protocol and sequenced on an Illumina MiSeq machine.

### tRNA sequence analysis

Illumina reads from tRNA-seq and RNA sequencing were mapped to the library of *E. coli* tRNAs, in which anticodons were localized (Supplementary Dataset 3). We used the Geneious mapper and Bowtie2 with a medium sensitivity within Geneious Prime (versions 2022-2024) for mapping of raw sequences to tRNAs. We identified mutations in anticodons and quantified the ratio of mutant to wild-type sequences for each tRNA. Raw tRNA and RNA sequences are provided under the NCBI bioproject ID defined in the Data Availability section.

### Purification of aminoacyl-tRNA synthetases

*E. coli* seryl-tRNA synthetase was PCR amplified from the genome of *E. coli* K12 MG1655 and cloned with a histidine tag into a ColE1 backbone with *bla* resistance marker and transformed into *E. coli* BL21(DE3) expression strain. The seryl-tRNA synthetase was overexpressed in *E. coli* cells until an OD_600_ of 0.5. Protein production was induced by adding 0.5 mM isopropyl-b-D-1-thiogalactopyranoside (Millipore Sigma) and cells were moved to a 33°C incubator and grown at 250 rpm for 2.5 hours. Cells were harvested at 6,000×g for 15 minutes at 4°C, washed with 1X PBS buffer, and stored at -80°C. The frozen cell pellet was thawed in lysis buffer (100 mM HEPES pH 7.2, 500 mM NaCl, 5mM BME) The cell paste was suspended in 15 mL of lysis buffer (50 mM Tris (pH 7.5), 300 mM NaCl, 20 mM imidazole) and lysed by sonication. The crude extract was centrifuged at 30,000×g for 30 minutes at 4°C. The soluble fraction was loaded onto a column containing 2 mL of Ni-NTA resin (Qiagen) previously equilibrated with 20 mL lysis buffer. The column was washed with a 20 mL lysis buffer, and the bound protein was then eluted with 2 mL of 50 mM Tris (pH 7.5), 300 mM NaCl, 300 mM imidazole. Purified proteins were dialyzed with 10 mM Tris (pH 7.5), 50, 1 mM DTT and 50% glycerol, and stored at −80°C for further studies.

### Production of tRNA *in vitro*

Oligonucleotides containing tRNA under a T7 promoter were synthesized by IDT and dsDNA was amplified by PCR. In vitro transcription (IVT) reactions were carried out at 37°C using the amplified tRNA dsDNA as template. IVT tRNA was purified and size selected using SPRIselect beads. The tRNA was refolded by heating to 90°C for 5 minutes and slow cooling to room temperature. At 65°C, MgCl_2_ was added to a final concentration of 10 mM.

### Aminoacylation activity assay

A 20 μL aminoacylation reaction contained the following components: 50 mM Tris-HCl (pH 7.2), 10 mM MgCl2, 10 mM ATP, 2 mM amino acids, 500 nM seryl-tRNA synthetase, and 20μg tRNA. Aminoacylation reactions were incubated at one hour at 37°C. The reactions were stopped by addition 5 μL of digestion solution consisting of 200 mM sodium acetate (pH5.2) and 1.5 U/μL RNAse A and incubated for 20 minutes at room temperature. Protein was precipitated by addition of 1% formic acid, and reactions were then frozen at -80oC from 30 minutes to 1 day. After precipitating protein at −80°C for 30 minutes, insoluble material was removed by centrifugation at 16,000×g for 15 minutes at 4°C. The soluble fraction was then transferred to autosampler vials, kept on ice until immediately before high resolution LC–MS analysis and returned to ice immediately afterwards.

### Mass spectrometry analysis of tRNA aminoacylation

Quantitative mass spectrometry data was collected using an Agilent 6530 Quadrupole Time-Of-Flight (QTOF) MS with an electrospray ionization (ESI) source, coupled to an Agilent Infinity 1290 ultra-high-performance liquid chromatography (UHPLC) system with an Agilent Poroshell 120 EC-C 18 2.7 μm, 2.1 x 50 mm column. Solvents used were (solvent A) water and 0.1% formic acid and (solvent B) acetonitrile and 0.1% formic acid. Mass spectra were gathered using Dual Agilent Jet Stream (AJS) ESI in positive mode. The mass range was set from 100 to 1700 m/z with a scan speed of 3 scan/second. The capillary and nozzle voltages were set to 3500 and 1000 V, respectively. The source parameters were set with a gas temperature of 325°C and a flowrate of 12 L/min, nebulizer at 35 psi, and sheath gas temperature at 350°C at a flow of 11 L/min. MS data were acquired with MassHunter Workstation Data Acquisition (Version B.06.01, Agilent Technologies) and analyzed using MassHunter Qualitative Analysis (Version 10.0, Agilent Technologies).

## Data availability

Sequences of plasmids and genes designed in this study are available in Supplementary tables 2, 8, and 9. Plasmid maps are available in Supplementary Datasets 1 and 2. Sequences of genomes reannotated in this study like the genome of *E. coli* MDS42 or the genome of Ec_Syn61Δ3 with marked anticodons, TCR, and TAG codons are available in the Supplementary Dataset 3. Plasmids used in this study are available at Genscript Inc. with IDs defined in Supplementary tables. The updated recoded strain Ec_Syn61Δ3 with genetic-code-sensitive kill switches and reporter generated in this study are available from Addgene (bacterial strains no. XXX and XXX). We have deposited raw data from whole-genome sequencing, plasmid sequencing, small RNA sequencing, and tRNA-seq to NCBI SRA BioProject under the ID PRJNA1136965, and raw data from proteomics at MassIVE under the ID MSV000095085_reviewer.

## Acknowledgements

This work was funded by the U.S. Department of Energy, Office of Science, Office of Biological and Environmental Research, under Award DE-FG02-02ER63445, the National Science Foundation, under Award 2123243 (both to G.M.C.), and Internal funding from the Wyss Institute. A.C-P. was also funded by the Swiss National Science Foundation, under Award P2ELP2_181884 (to A.C-P.).

We thank Ryan Rogers-Hammond and Raphaël Ferreira for feedback on the manuscript; Suyay Chiappino Pepe for help with Supplementary Fig. 1; Gregory Stephanopoulos and Aarti Krishnan for general discussions on the project; George Chao for discussions on DNA sequencing and image analysis; Max Schubert for discussions on library screening; Russel M Vincent for discussions on tRNA modifications; Svenja Vinke for discussions on non-standard amino acids; Akos Nyerges for sharing the strain Ec_Syn61Δ3-SL; the rest of the Church lab members for general discussions and support. The first author would also like to thank Florian Wehner and Hugo Wehner for their love and support.

We acknowledge the companies SeqCenter (prev. MIGS) for Illumina sequencing of genomic DNA samples and tRNA-seq samples; Novogene for small RNA sequencing; Azenta and Quintara Bio for next generation sequencing of bipARS libraries; Azenta for Sanger sequencing of amplicons and primer synthesis; IDT for DNA oligo and gBlock synthesis; Plasmidsaurus for nanopore plasmid sequencing; and Genscript U.S.A. Inc. for DNA and plasmid synthesis.

## Author contributions

Project administration: G.M.C., A.C-P., J.M.T.; Conceptualization: G.M.C., A.C-P.; Resources: G.M.C., A.C-P., J.M.T., B.B.; Data curation: A.C-P., F.R., B.B., T.L.A.; Software: A.C-P., M.M., B.B., T.L.A.; Formal analysis: A.C-P., F.R., B.B., T.L.A., M.M.; Investigation: A.C-P.; Visualization: A.C-P., B.B., M.M., A.M.D.; Methodology: G.M.C., A.C-P., F.R., B.B., H.T.; Writing – original draft: A.C-P.; Writing – review and editing: all authors; Funding acquisition: G.M.C., A.C-P., J.M.T., K.N., J.D.A.; Experimental discovery of escapees: A.C-P.; Design of genes and plasmids: A.C-P.; Acquisition of samples: A.C-P.; Analysis of sequencing and omics data: A.C.-P., B.B., T.L.A.; Proteomics: B.B.; Evaluation of bipARS variants: T.L.A., A.C-P.; tRNA-seq: F.R., H.M.B.; *in vitro* tRNA studies: F.R.; DNA synthesis: Q.Z., W.F.; Preparation of LB agar plates: M.M.A., S.T., A.C-P.; Evaluation of reporters: A.C-P., A.M.D., T.L.A.; Computational analysis of antibacterials: A.C-P., M.M.; Contributing to protocol definition, sharing reagents: G.M.C., A.C-P., F.R., B.B., H.T., E.K., K.N., J.A.M., R.P., J.D.A., J.M.S.; Bioethical considerations: J.E.L.

