## Supplementary Fig. for "Preventing escape and malfunction of recoded cells due to tRNA base changes"

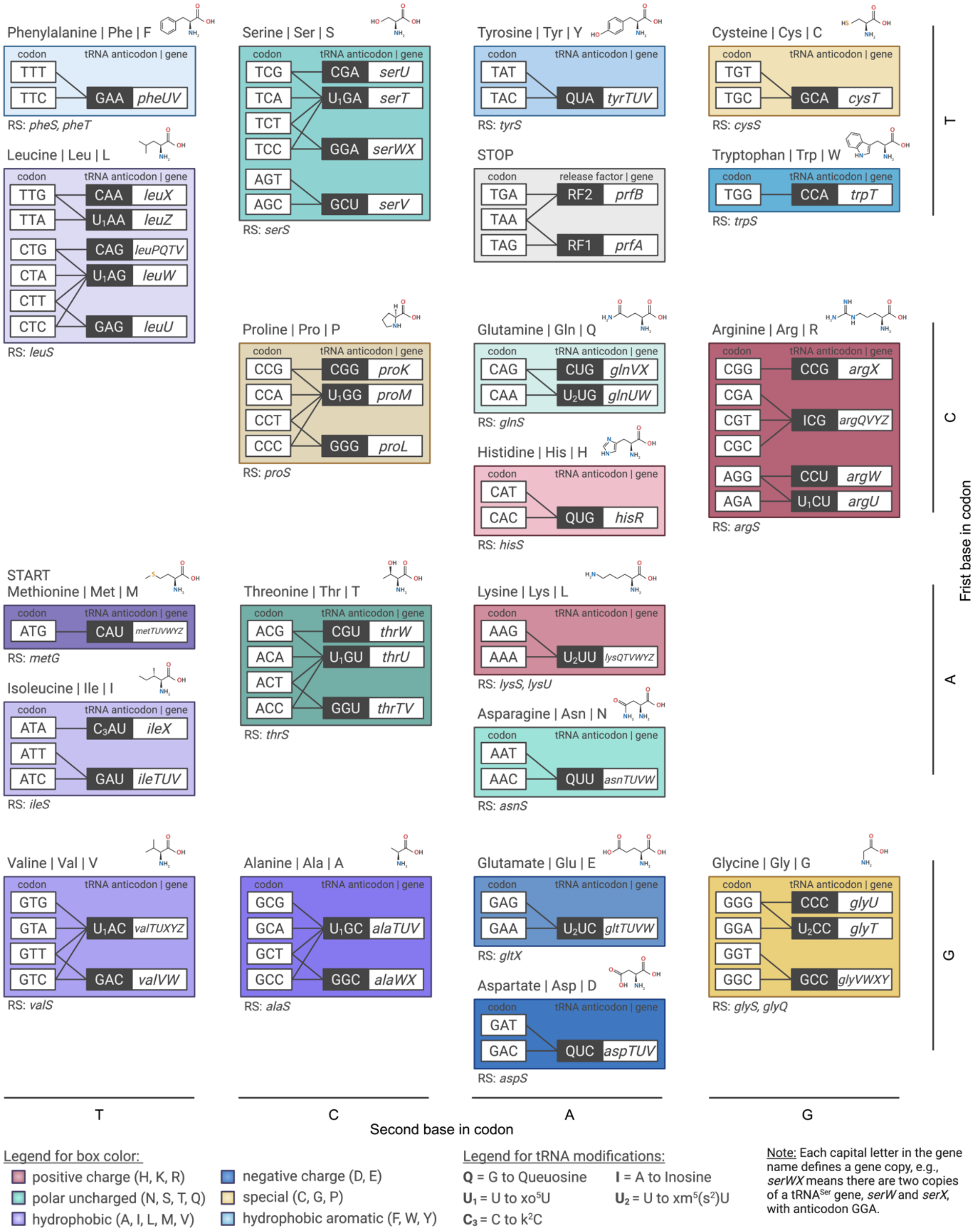


**Supplementary Fig. 1. The standard genetic code in *Escherichia coli*.** We include all known interactions between codons, tRNAs, and aminoacyl-tRNA synthetases (RS, below boxes) in *Escherichia coli*. This information applies to K-12 MG1655 and MDS42 *E. coli* strains, which are parental strains of Ec_Syn61Δ3. We group codons based on the amino acid they code for. We group codons, to the extent possible, based on the first, second, and third base. We link amino acids considering properties like charge and hydrophobicity based on box colors, as defined in the legend. We define known tRNA anticodon modifications. Each capital letter in the gene name of a tRNA indicates a gene copy, *e.g.*, *serWX* indicates there are two copies *serW* and *serX*.

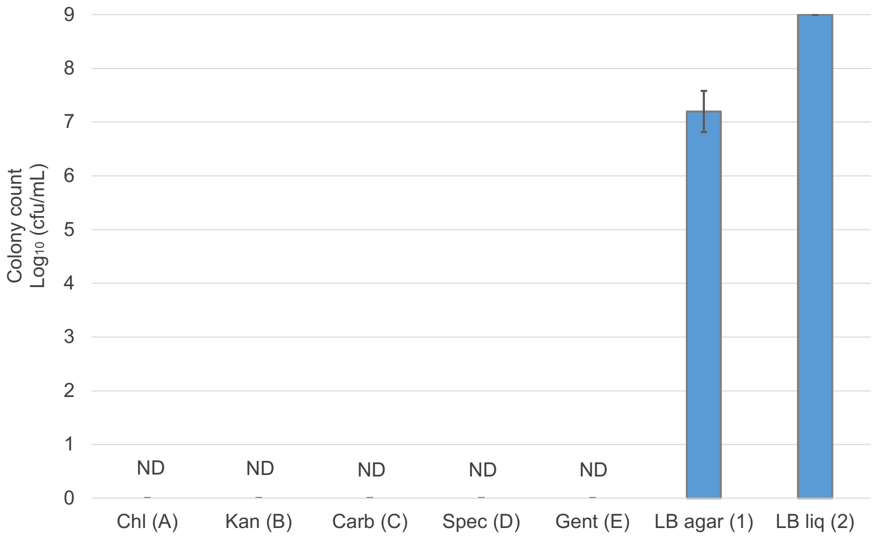


**Supplementary Fig. 2. Escape rate of plasmid-less Ec_Syn61Δ3.** Colonies formed by plasmid-less Ec_Syn61Δ3 onto agar plates containing standard antibiotic concentrations (*n* = *5*). ND denotes *none detected*. Results from two tests. In test 1, plasmid-less Ec_Syn61Δ3 underwent electroporation, overnight growth, plating, and selection onto agar plates containing standard antibiotic concentrations (*n* = *2*), following the same approach as experiments in which escapees were observed (Fig. 1). The LB agar (1) bar shows the number of colonies formed by these same cultures in a Luria Broth (LB) agar plate without antibiotics (mean ± SD, *n* = *10*). In test 2, plasmid-less Ec_Syn61Δ3 did not undergo electroporation; it was grown overnight, plated, and selected onto antibiotic-containing agar plates (*n* = *3*). The LB liq (2) bar shows the number of cells plated based on a normalized $OD_{600}$ = 1 (*n* = *15*). Antibiotic-containing agar plates were incubated for 10 days at 37°C, and LB plates were incubated for 3 days at 37°C. Antibiotic acronyms: Chl (A), chloramphenicol; Kan (B), kanamycin; Carb (C), carbenicillin; Spec (D), spectinomycin; and Gent (E), gentamicin.

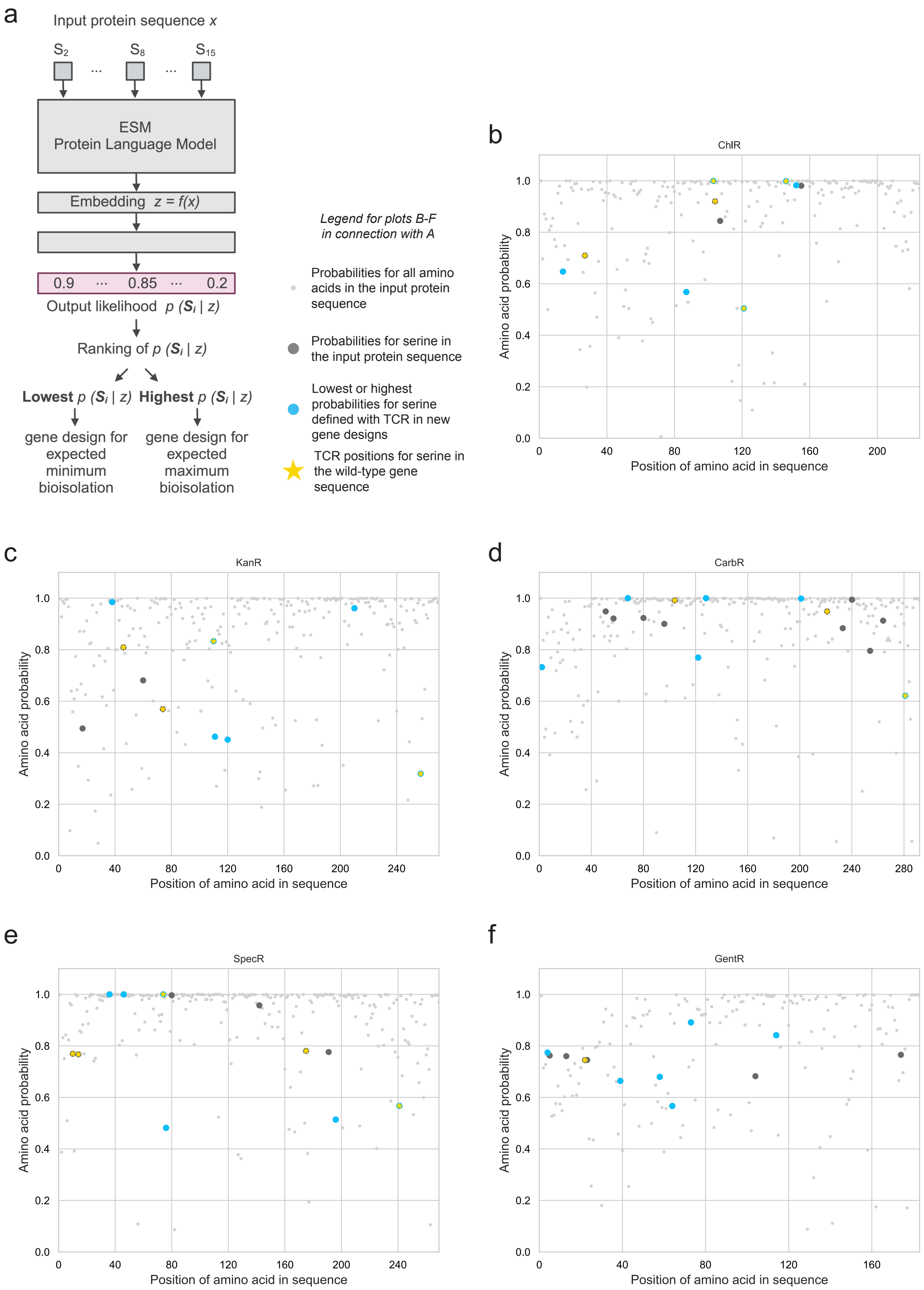


**Supplementary Fig. 3. Design of genes with a protein language model. a,** Pipeline used to design genes with TCR codons in positions with the maximum and minimum probability to incorporate serine based on a protein language model (PLM) ESM. **b-f,** Computed amino acid probabilities at each position in the input protein sequence based on the PLM ESM for antibiotic resistance genes. We show four groups of data: 1) probabilities for all amino acids (small light gray dots); 2) probabilities for all serine positions (large dark gray dots, or blue if they overlap with group three); 3) top and bottom probabilities for serine, which are TCR positions in our gene designs (large blue dots); 4) probabilities for serine in TCR positions of wild-type gene designs (overlapping yellow star). Antibiotic resistance acronyms: ChlR, chloramphenicol; KanR, kanamycin; CarbR, carbenicillin; SpecR, spectinomycin; and GentR, gentamicin.

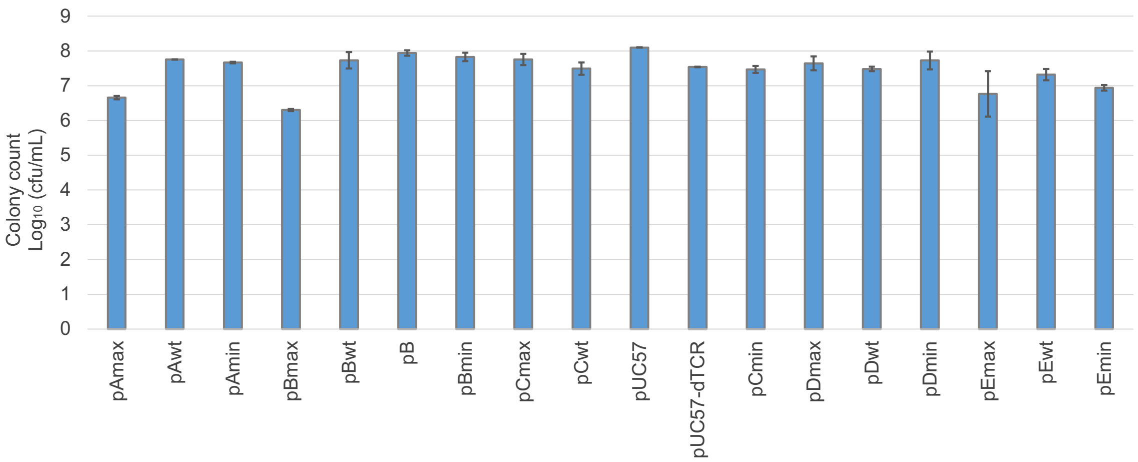


| | | | | | | | | | | | | | | | | | | |
|---|---|---|---|---|---|---|---|---|---|---|---|---|---|---|---|---|---|---|
| TCR count | 3 | 5 | 3 | 3 | 4 | 4 | 3 | 3 | 3 | 3 | 3 | 3 | 3 | 5 | 3 | 3 | 1 | 3 |
| Plasmid copy | + | + | + | + | + | + | + | + | + | + | + | + | + | + | + | + | + | + |
| His-tag | + | + | + | + | + | - | + | + | + | - | - | + | + | + | + | + | + | + |
| Antibiotic resistance | ChlR | | | KanR | | | | AmpR/CarbR | | | | | SpecR | | | GentR | | |

**Supplementary Fig. 4. Demonstration of functionality of newly designed genes.** Colonies formed by MDS42ΔrecA, a reference *E. coli* strain with *serT* and *serU* $tRNA^{Ser}$ genes and parental strain of Ec_Syn61Δ3, after transformation with plasmids designed in this study, including two available plasmids pB and pUC57, for reference (mean ± SD; $n$ = 2). Results show expression and functionality of the designed genes in the presence of $tRNA^{Ser}$ genes *serT* and *serU*. We followed the same protocols for competent cell preparation, electroporation, plating, and selection as done for Ec_Syn61Δ3 (Fig. 1). Table below describes number of TCR codons, plasmid copy number, presence of a polyhistidine tag on the antibiotic resistance protein, and antibiotic resistance for each plasmid. High or low plasmid copy number is denoted by + or -. Presence or absence of a His-tag is denoted by + or -. Antibiotic resistance acronyms: ChlR, chloramphenicol; KanR, kanamycin; AmpR/CarbR, ampicillin/carbenicillin; SpecR, spectinomycin; and GentR, gentamicin.

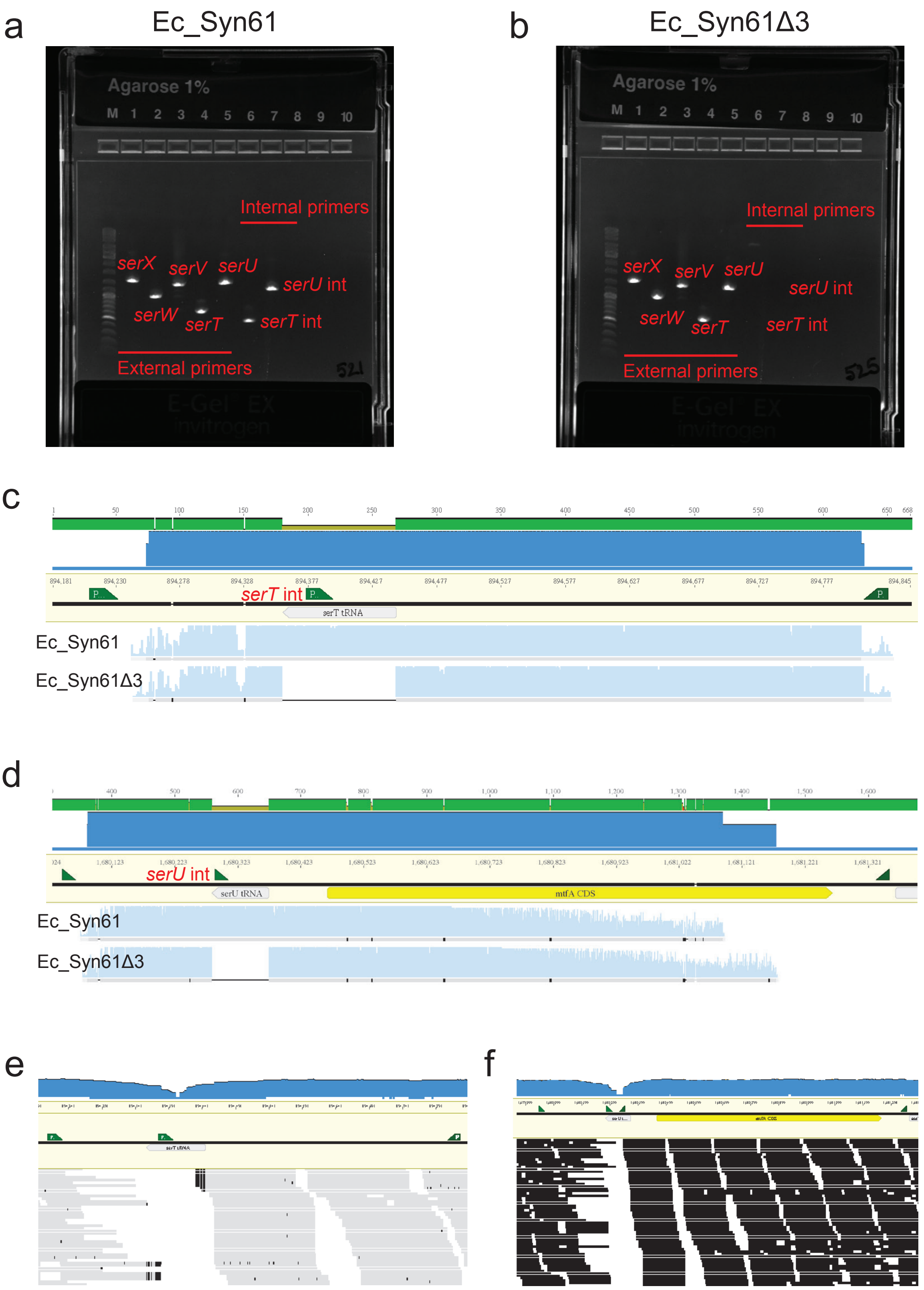


**Supplementary Fig. 5. Verification of *serT* and *serU* genes absence in Ec_Syn61Δ3. a,** Gel image of amplicons for verification of Ec_Syn61 genomic *loci* sequence around tRNA$^{Ser}$ genes *serX* (band 1), *serW* (band2), *serV* (band 3), *serT* (band 4) and *serU* (band 5) using external primers. We verified the presence of *serT* (band 6) and *serU* (band 7) genes in Ec_Syn61 with internal primers. **b,** Gel image of amplicons for verification of Ec_Syn61Δ3 genomic *loci* sequence around tRNA$^{Ser}$ genes *serX* (band 1), *serW* (band2) and *serV* (band 3), including loci where *serT* (band 4) and *serU* (band 5) genes where previously located, using external primers. We verified the absence of tRNA$^{Ser}$ genes *serT* (band 6) and *serU* (band 7) in Ec_Syn61Δ3 with internal primers. **c,** Sanger sequencing of tRNA$^{Ser}$ *serT* loci amplicons from Ec_Syn61 (first row of reads) and Ec_Syn61Δ3 (second row of reads) genomes, using MDS42 as the reference genome map sequence. **d,** Sanger sequencing of tRNA$^{Ser}$ *serU* loci amplicons from Ec_Syn61 (first row of reads) and Ec_Syn61Δ3 (second row of reads) genomes, using MDS42 as the reference genome map sequence. **e,** Illumina sequencing of Ec_Syn61Δ3 genome mapped for reference to the *E. coli* strain MDS42 genome map to verify absence of tRNA$^{Ser}$ gene *serT*. **f,** Illumina sequencing of Ec_Syn61Δ3 genome mapped for reference to the *E. coli* strain MDS42 genome map to verify absence of tRNA$^{Ser}$ gene *serU*.

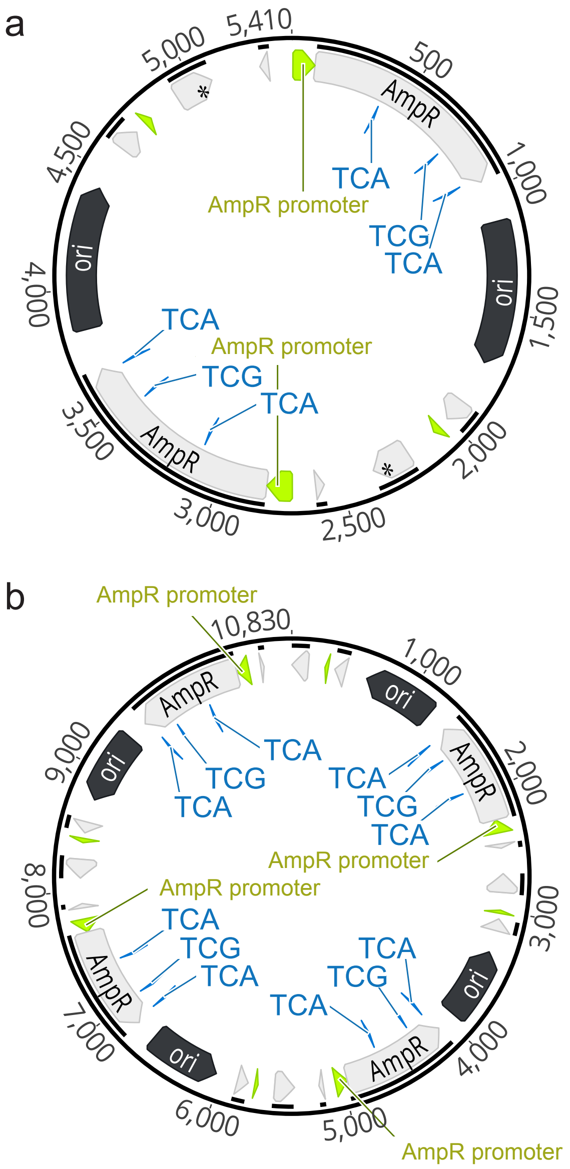


**Supplementary Fig. 6. Example sequencing results of a plasmid in Ec_Syn61Δ3 escapees.** This example plasmid pUC57 shows two (a) and four (b) copies of the carbenicillin resistance gene coding for beta-lactamase based on long read sequencing. All gene copies keep all three initial instances of TCR codons. Raw sequences are available under the NCBI bioproject ID provided in the main manuscript.

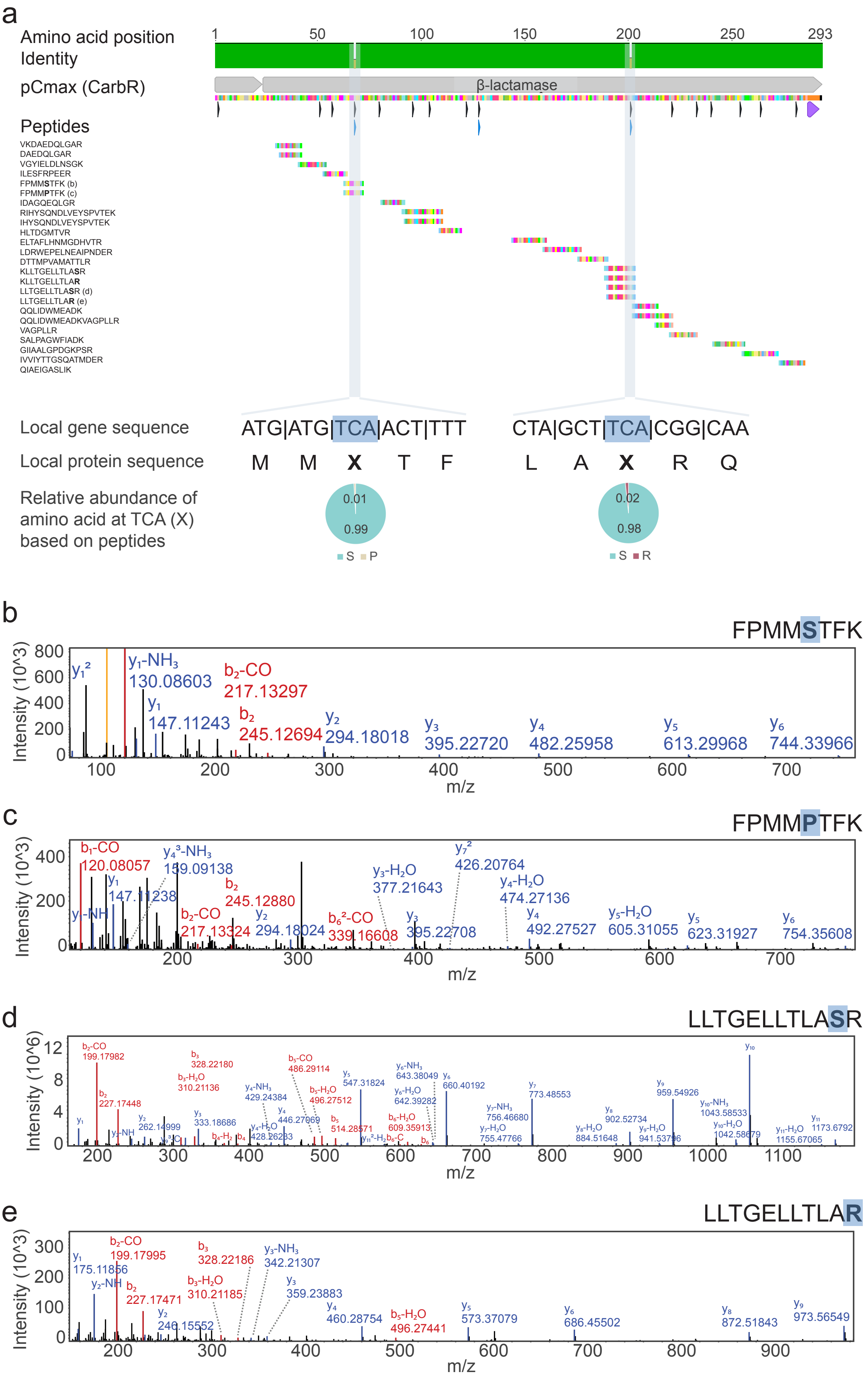
a
Amino acid position
Identity
pCmax (CarbR)
β-lactamase
Peptides
VKDAEDQLGAR
DAEDQLGAR
VGYIELDLNSGK
ILESFRPEER
FPMMSTFK (b)
FPMMPTFK (c)
IDAGQEQLGR
RIHYSQNDLVEYSPVTEK
IHYSQNDLVEYSPVTEK
HLTDGMTVR
ELTAFLHNMGDHVTR
LDRWEPELNEAIPNDER
DTTMPVAMATTLR
KLLTGELLTLASR
KLLTGELLTLAR
LLTGELLTLASR (d)
LLTGELLTLAR (e)
QQLIDWMEADK
QQLIDWMEADKVAGPLLR
VAGPLLR
SALPAGWFIADK
GIIAALGPDGKPSR
IVVIYTTGSQATMDER
QIAEIGASLIK
Local gene sequence
ATG|ATG|TCA|ACT|TTT
CTA|GCT|TCA|CGG|CAA
Local protein sequence
M M X T F
L A X R Q
Relative abundance of amino acid at TCA (X) based on peptides
0.01
0.99
S P
0.02
0.98
S R
b
FPMMSTFK
Intensity (10^3)
m/z
c
FPMMPTFK
Intensity (10^3)
m/z
d
LLTGELLTLASR
Intensity (10^6)
m/z
e
LLTGELLTLAR
Intensity (10^3)
m/z

**Supplementary Fig. 7.** Proteomics results showing beta-lactamase detection (conferring carbenicillin resistance) from expression of the pCmax gene design in Ec_Syn61Δ3 escapees (*n* = 2). **a,** Top: detected peptides across the beta-lactamase protein sequence, with highlights on peptides identifying different amino acids at TCR positions. Bottom: relative abundance of amino acids at TCR positions based on peptides' quantification. **b,** Mass spectrum showing detection of beta-lactamase with serine in the first of three TCR positions in the pCmax design with peptide FPMMSTFK. **c,** Mass spectrum showing detection of beta-lactamase with proline in the first of three TCR positions in the pCmax design with peptide FPMMPTFK. **d,** Mass spectrum showing detection of beta-lactamase with serine in the third of three TCR positions in the pCmax design with peptide LLTGELLTLASR. **e,** Mass spectrum showing detection of beta-lactamase with arginine in the third of three TCR positions in the pCmax design with peptide LLTGELLTLAR.

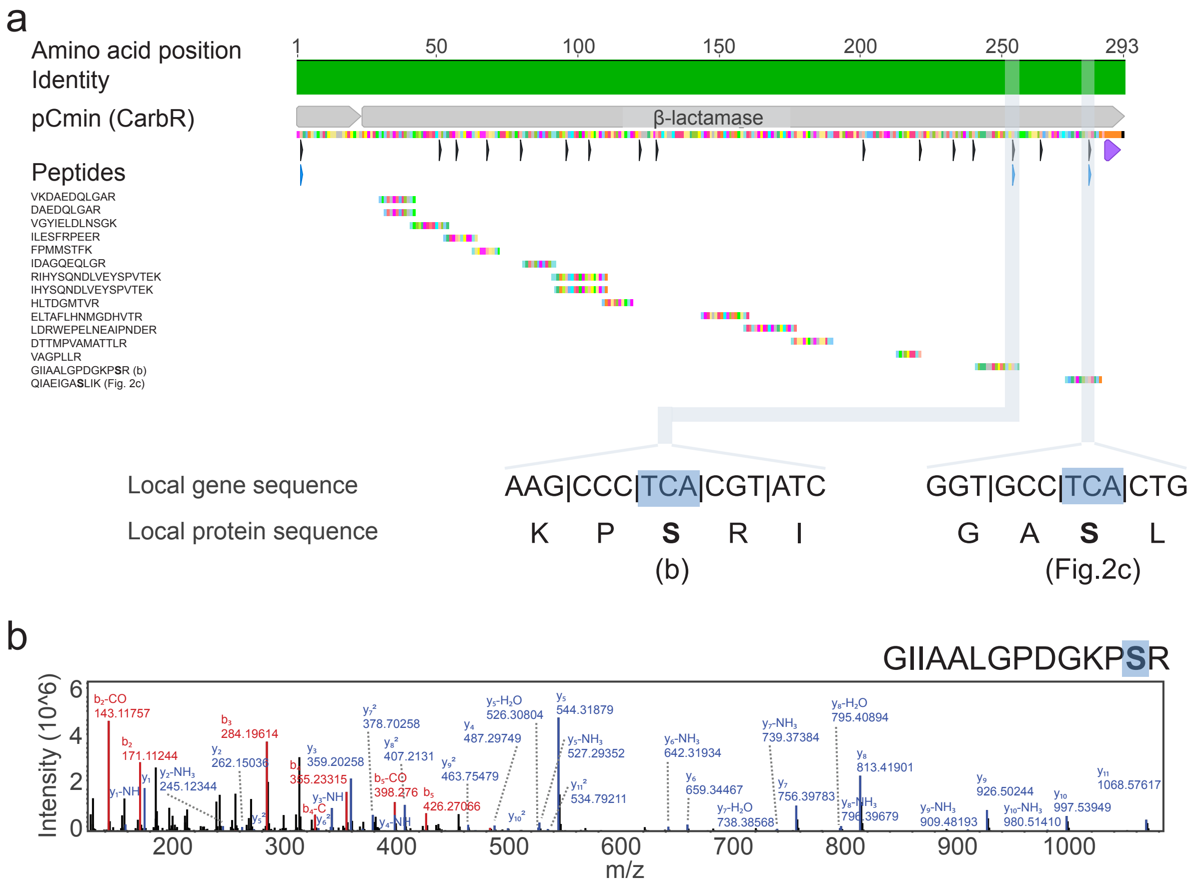


**Supplementary Fig. 8.** Proteomics results showing beta-lactamase detection (conferring carbenicillin resistance) from expression of the pCmin gene design in Ec_Syn61Δ3 escapees ($n$ = 3). **a,** Top: detected peptides across the beta-lactamase protein sequence, with highlights on peptides identifying serine at TCR positions. **b,** Mass spectrum showing detection of beta-lactamase with serine in the second of three TCR positions in the pCmin design with peptide GIIAALGPDGKPSR.

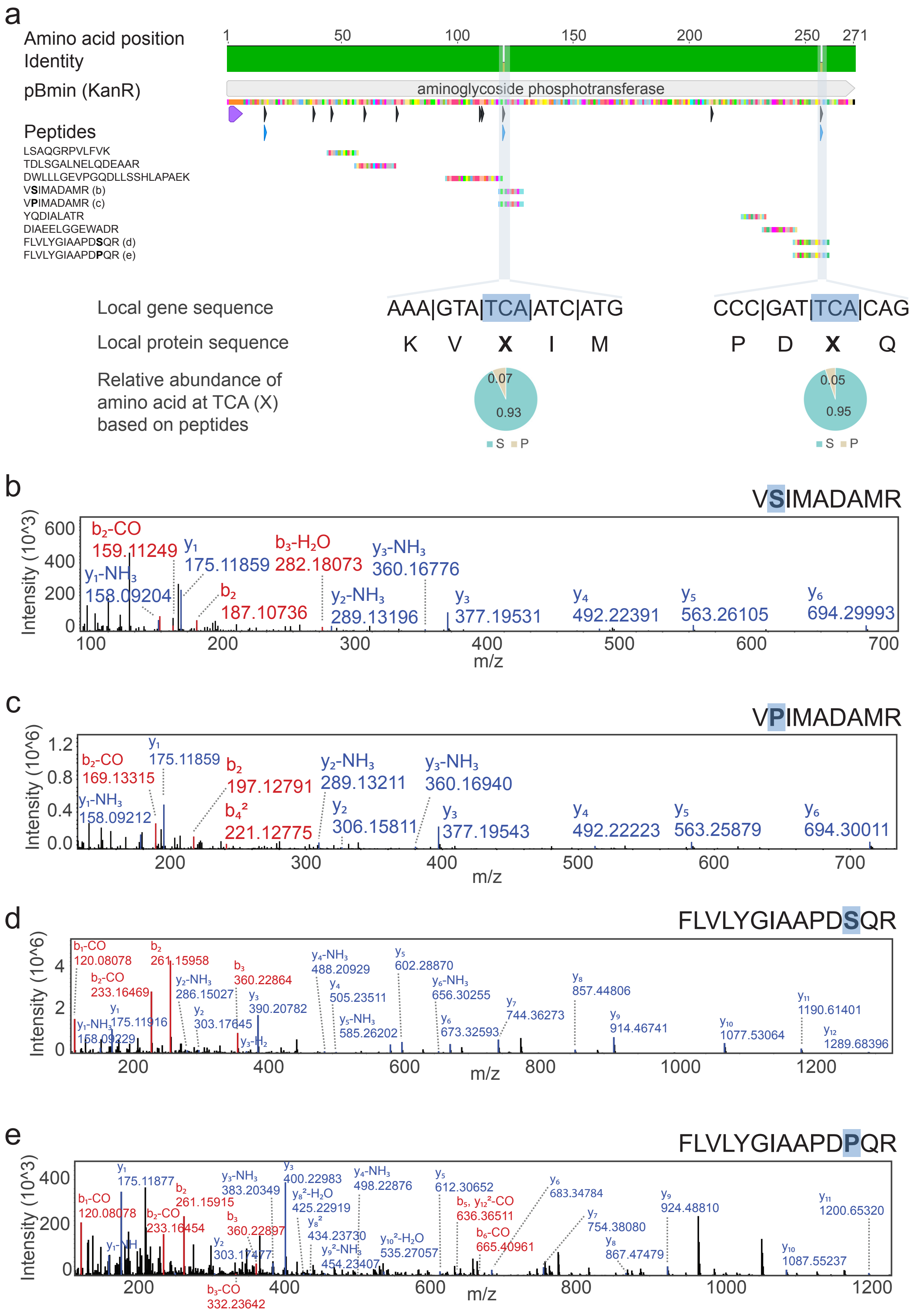


**Supplementary Fig. 9**. Proteomics results showing aminoglycoside phosphotransferase detection (conferring kanamycin resistance) from expression of the pBmin gene design in Ec_Syn61Δ3 escapees (*n* = 2). **a,** Top: detected peptides across the aminoglycoside phosphotransferase protein sequence, with highlights on peptides identifying different amino acids at TCR positions. Bottom: relative abundance of amino acids at TCR positions based on peptides' quantification. **b,** Mass spectrum showing detection of aminoglycoside phosphotransferase with serine in the second of three TCR positions in the pBmin design with peptide VSIMADAMR. **c,** Mass spectrum showing detection of aminoglycoside phosphotransferase with proline in the second of three TCR positions in the pBmin design with peptide VPIMADAMR. **d,** Mass spectrum showing detection of aminoglycoside phosphotransferase with serine in the third of three TCR positions in the pBmin design with peptide FLVLYGIAAPDSQR. **e,** Mass spectrum showing detection of aminoglycoside phosphotransferase with proline in the third of three TCR positions in the pBmin design with peptide FLVLYGIAAPDPQR.

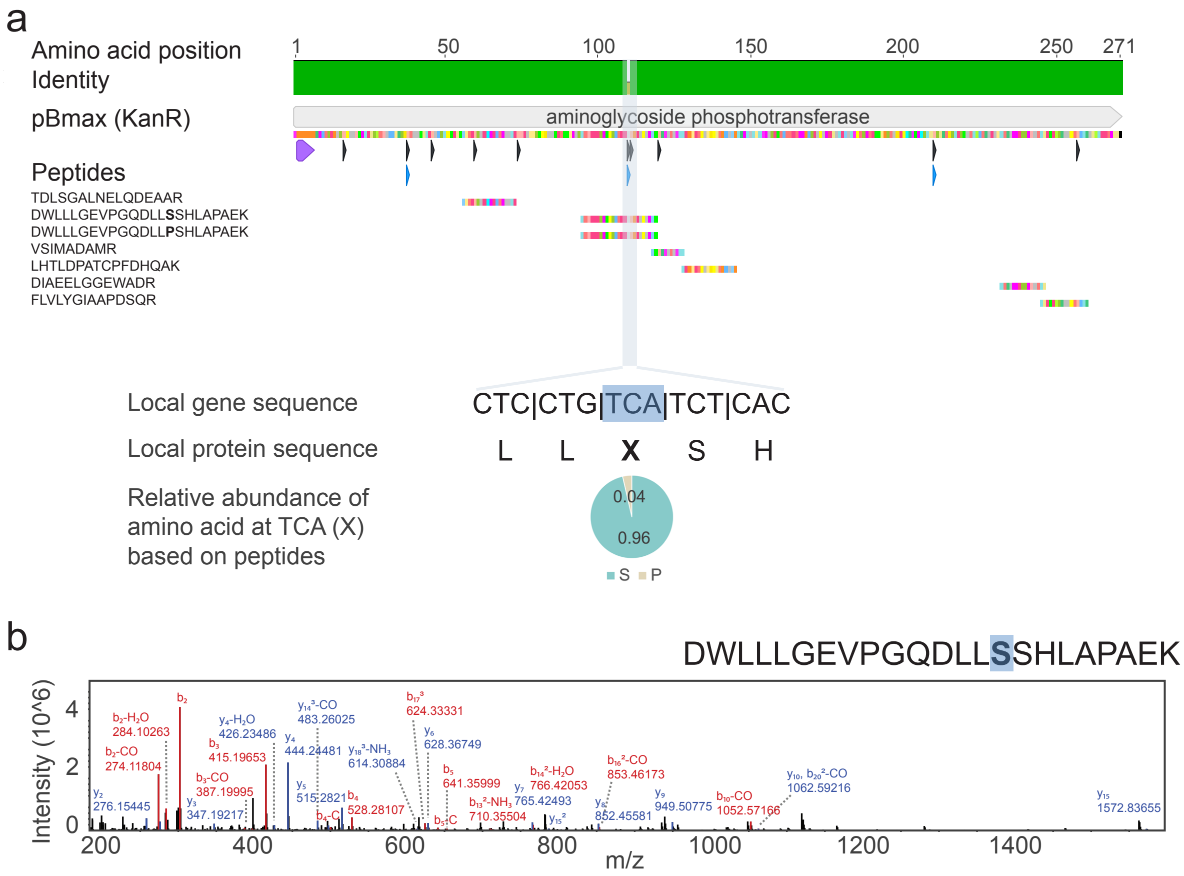


**Supplementary Fig. 10**. Proteomics results showing aminoglycoside phosphotransferase detection (conferring kanamycin resistance) from expression of the pBmax gene design in Ec_Syn61Δ3 escapees ($n$ = 2). **a,** Top: detected peptides across the aminoglycoside phosphotransferase protein sequence, with highlights on peptides identifying different amino acids at TCR positions. Bottom: relative abundance of amino acids at TCR positions based on peptides' quantification. **b,** Mass spectrum showing detection of aminoglycoside phosphotransferase with serine in the second of three TCR positions in the pBmax design with peptide DWLLLGEVPGQDLLSSRLAPAEK.

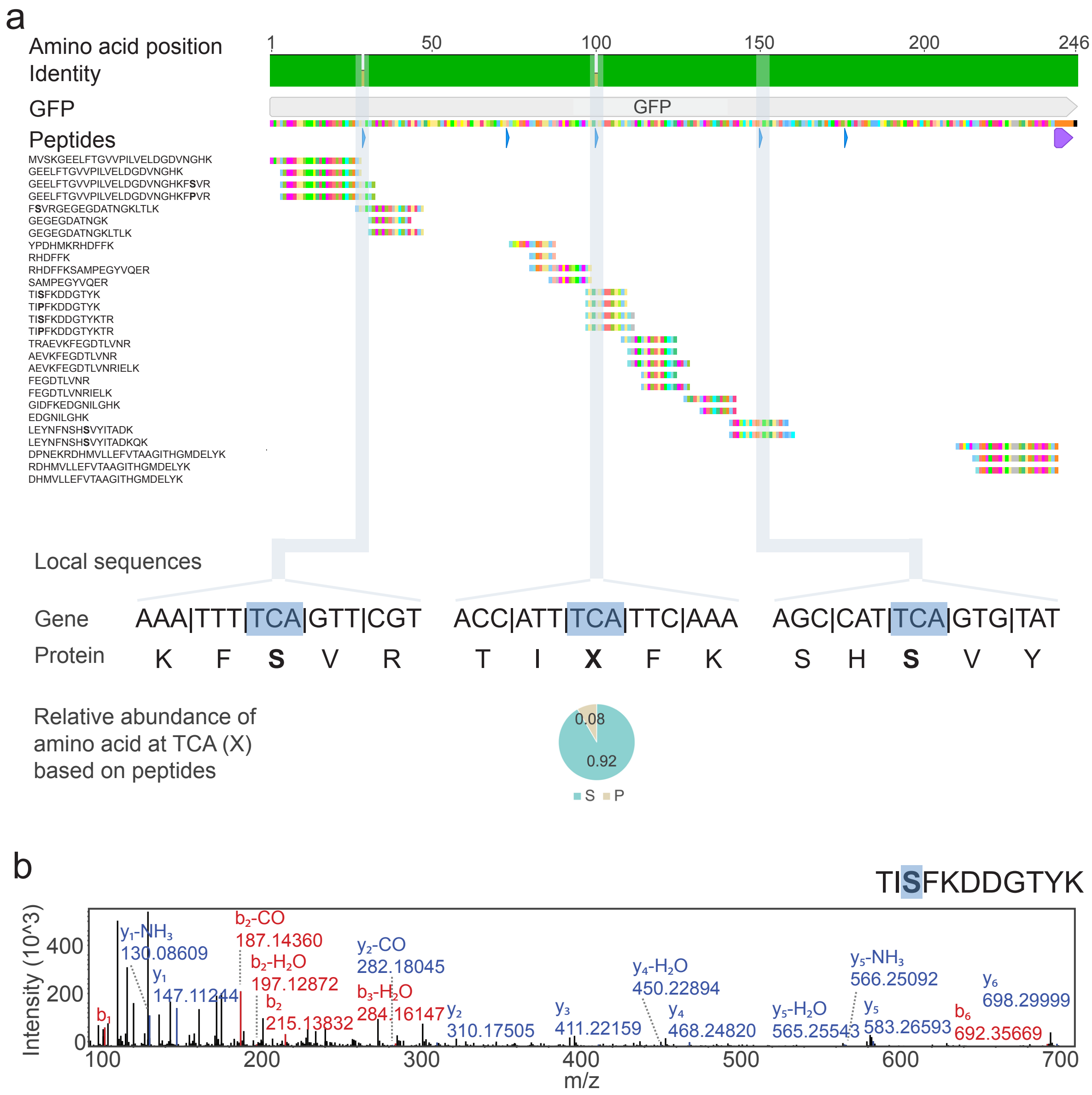


**Supplementary Fig. 11**. Proteomics results showing GFP detection from expression of our genetic-code-sensitive reporter gene design in Ec_Syn61Δ3 escapees (*n* = *9*). **a,** Top: detected peptides across the GFP sequence, with highlights on peptides identifying serine or different amino acids at TCR positions. Bottom: relative abundance of amino acids at TCR positions based on peptides' quantification. **b,** Mass spectrum showing detection of GFP with serine in the third of five TCR positions in the genetic-code-sensitive reporter gene design with peptide TISFKDDGTYK.

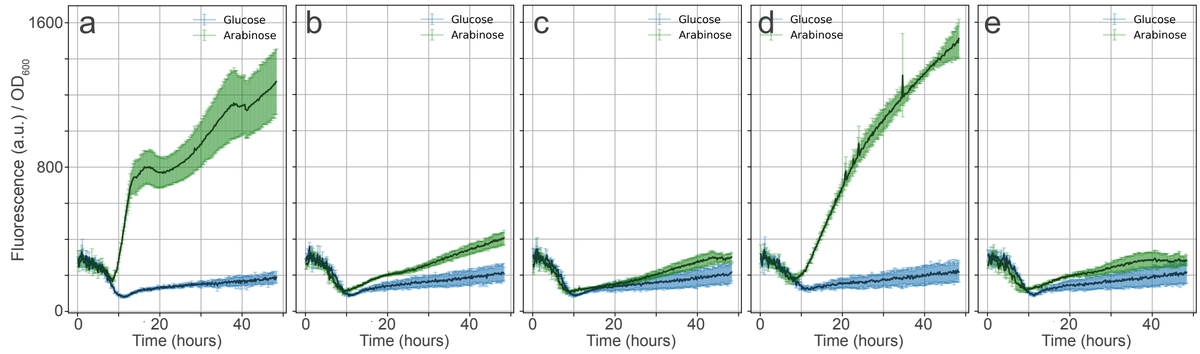


| | a | b | c | d | e |
|---|---|---|---|---|---|
| TCR count | 1 | 2 | 3 | 5 | 5 |
| TCR position | 150 | 2, 150 | 2, 3, 150 | 29, 73, 100, 150, 176 | 29, 73, 100, 150, 176 |
| Recoded AbxR gene | + | + | + | - | + |

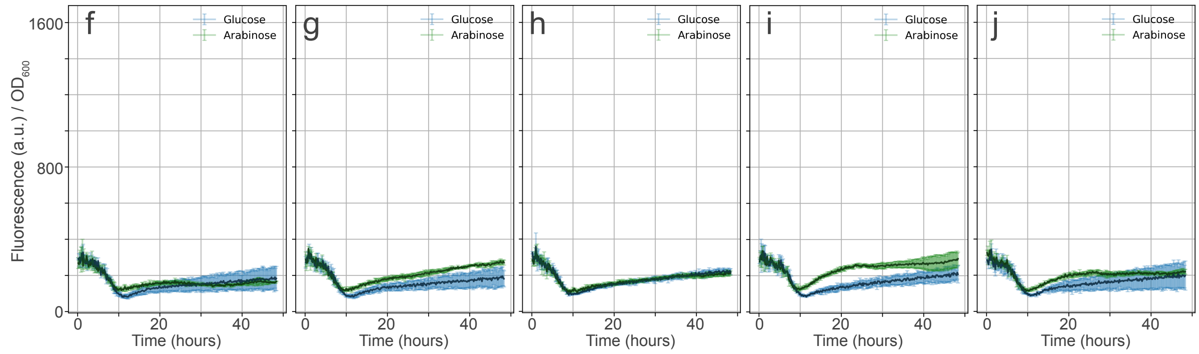


| | f | g | h | i | j |
|---|---|---|---|---|---|
| TCR count | 2 | 2 | 2 | 2 | 2 |
| TCR position | 182, 185 | 49, 219 | 71, 78 | 78, 79 | 79, 82 |
| Recoded AbxR gene | + | + | + | + | + |

**Supplementary Fig. 12**. Performance of alternative GFP reporters in Ec_Syn61Δ3. Fluorescence per $OD_{600}$ over time in Ec_Syn61Δ3 with alternative GFP reporters under an arabinose inducible promoter (mean ± SD, *n = 3*). GFP expression is induced with arabinose (green) and repressed with glucose (blue). Table below highlights number of TCR codons and positions of those TCR codons in each GFP design. All genes coding for GFP are part of a plasmid with a recoded carbenicillin resistance gene or pUC57-dTCR, except for (d) which shows an Ec_Syn61Δ3 escapee with a wild-type carbenicillin resistance gene.

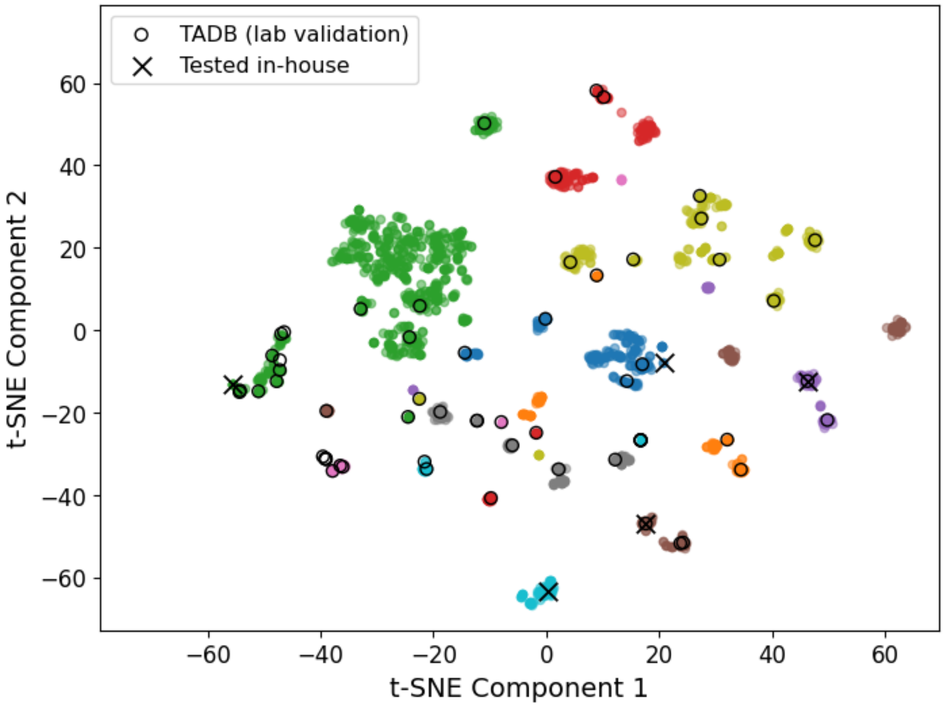


**Supplementary Fig. 13. Analysis of antibacterials or toxins for design of our genetic-code-sensitive kill switches**. t-SNE projection of the antibacterial or toxin peptide sequence. Colored dots: toxins from the database TADB 3.0, colored according to a DBSCAN clustering. Black circles: toxins from the experimental validation subset of TADB 3.0. Black X: toxins tested in house with two gene designs.

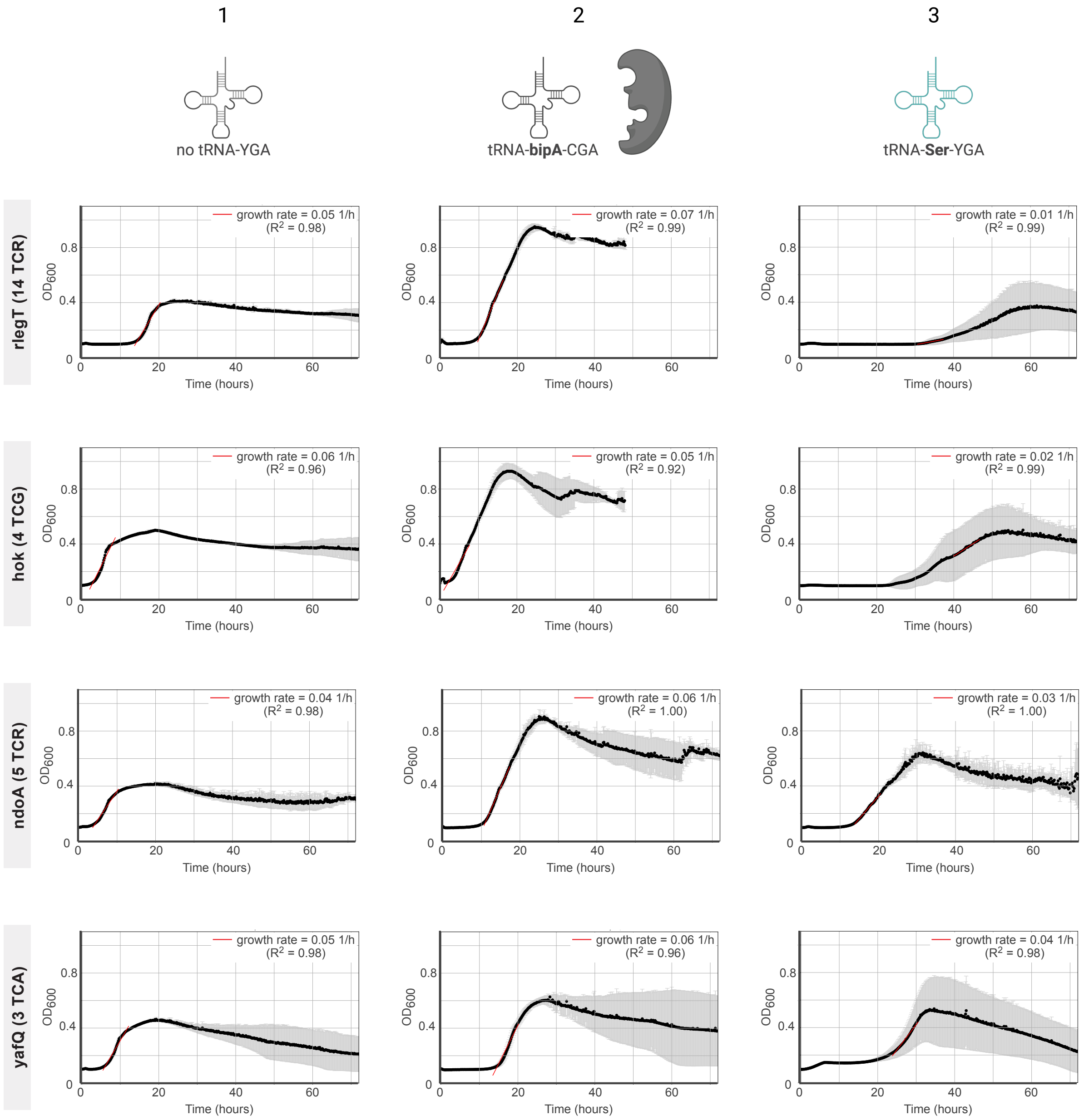


**Supplementary Fig. 14**. Performance of our genetic-code-sensitive kill switches. We show four alternative designs (rlegT$_{14\ TCR}$, hok$_{4\ TCG}$, ndoA$_{5\ TCR}$, yafQ$_{3\ TCA}$, Supplementary table 8) that do not significantly delay growth or affect growth rate when there is no wild-type tRNA$^{Ser}$ cognate to TCR codons in Ec_Syn61Δ3 (column 1, mean ± SD, $n$ = 5) or when there is an orthogonal tRNA$^{bipA}_{CGA}$ and bipA aminoacyl-tRNA synthetase (bipARS) pair (column 2, mean ± SD, $n$ = 3). These designs also show different performance considering the growth delay and growth rate of Ec_Syn61Δ3 upon low level expression of tRNA$^{Ser}$ genes *serT* and *serU* (column 3, mean ± SD, $n$ = 5). Growth rates represent the slope of the red line fit to the curve, for which we provide the R-square value (Methods). All experiments started with a normalized number of ~$10^6$ cells at $OD_{600}$=0.01.

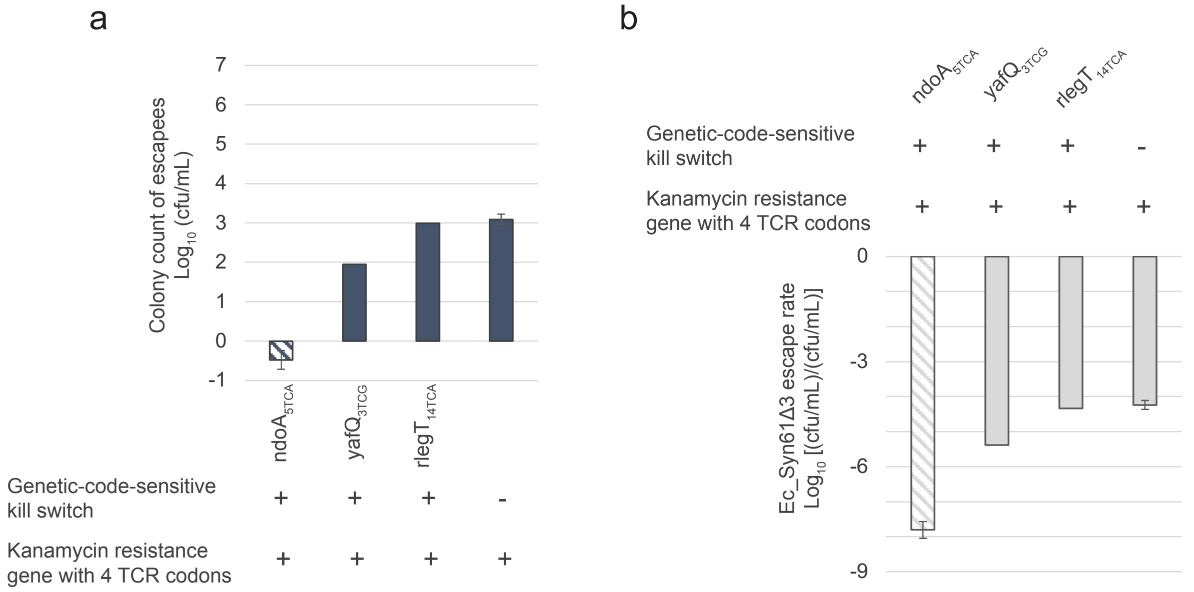


**Supplementary Fig. 15**. Escape rate of Ec_Syn61Δ3 with a set of genetic-code-sensitive kill switches in the presence of a kanamycin resistance gene with 4 TCR codons (pB design). Designs of genetic code sensitive kill switches include ndoA$_{5\ TCA}$, yafQ$_{3\ TCG}$, and rlegT$_{14\ TCA}$. **a,** Colony count of escapees for parallel experiments with varying kill switches. For ndoA$_{5\ TCA}$, we detected one escapee in a single replicate and no escapees in two additional replicates (first bar). The number of colonies of Ec_Syn61Δ3 without kill switch and the pB plasmid alone (last bar) was also shown in Fig 1d. **b,** Escape rates defined as the number of colonies found divided by the average number of colonies formed ($2.1x10^7$ cfu/mL) by plasmid-less Ec_Syn61Δ3 onto LB agar plates after electroporation in parallel experiments to those showing escapees.

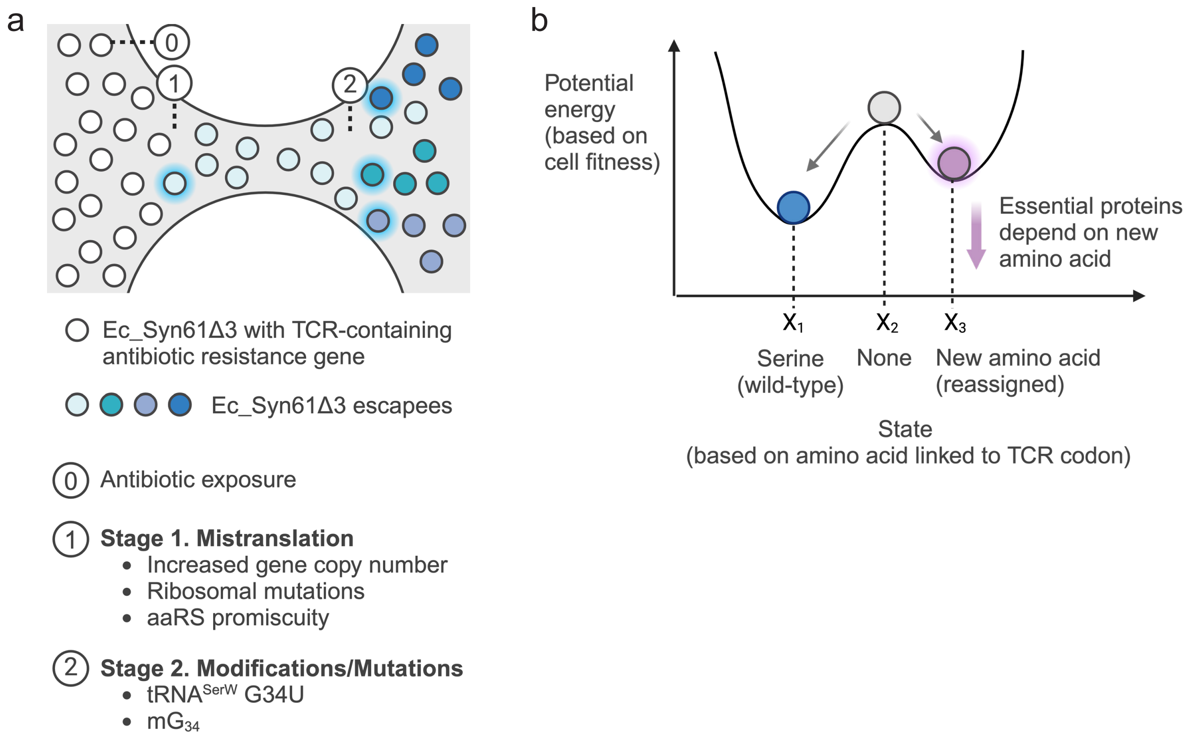


c

**Elements influencing likelihood of escape from genetic-code-based cellular bioisolation and malfunction due to misincorporation of standard amino acids (here, serine)**

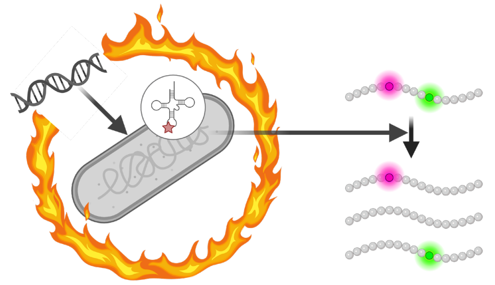

**Transferred DNA with TCR-containing genes**

*Elements determining gene expression rate and level*

| Element in transferred DNA | Design maximizing escape rate |
|---|---|
| Effect of gene on fitness | Essential / fitness advantage |
| Number of TCR codons | <5 |
| Position of TCR codons | Low serine probability |
| Polyhistidine tag (His-tag) | Absence |
| Promoter | Strong |
| Ribosome binding site (RBS) | Strong |
| Plasmid/gene copy number | High copy |

**Culture conditions**

*Context specific elements determining needs for survival*

| Element in context | Design maximizing escape rate |
|---|---|
| Antibiotic type | Low stringency |
| Antibiotic concentration | Low concentration |
| Rich/minimal medium | Rich |
| Incubation days | 3-8 days |
| Temperature | 37°C |
| Agitation/Aeration | Presence |
| Cell concentration | >$10^2$-$10^7$ cfu/mL |

**Cellular state**

*Compatibility of genetic code from incoming gene with genetic code of recoded cell*

| Element in cell | Design maximizing escape rate |
|---|---|
| % Genome recoded | <100% |
| Ribosome fidelity | Low |
| Promiscuous aaRSes | Presence |
| Nb. of aaRS/amino acid | >1 |
| tRNA copy number | >1 |
| tRNA modifying enzymes | Presence |
| "Free" codons | Not reassigned |

**Supplementary Fig. 16. Considerations for the design, applications, and biosafety of synthetic genomes and genes in recoded cells. a**, Conceptual representation of the discovered mechanism of escape in Ec_Syn61Δ3 acquiring antibiotic resistance through expression of TCR-containing antibiotic resistance genes. Upon antibiotic selection Ec_Syn61Δ3 acquires antibiotic resistance in a two-stage process involving mistranslation and mutations, modifications of essential macromolecules of the translation machinery. **b**, Conceptual representation of the stability of an engineered genetic code based on the recoded codon reassignment. **c**, Elements playing a role in the expression of TCR-containing genes in Ec_Syn61Δ3, which should be generalizable to recoded cells with other recoding schemes.
